# Targeted micro-fiber arrays for measuring and manipulating localized multi-scale neural dynamics over large, deep brain volumes during behavior

**DOI:** 10.1101/2023.11.17.567425

**Authors:** Mai-Anh T. Vu, Eleanor H. Brown, Michelle J. Wen, Christian A. Noggle, Zicheng Zhang, Kevin J. Monk, Safa Bouabid, Lydia Mroz, Benjamin M. Graham, Yizhou Zhuo, Yulong Li, Timothy M. Otchy, Lin Tian, Ian G. Davison, David A. Boas, Mark W. Howe

## Abstract

Neural population dynamics relevant for behavior vary over multiple spatial and temporal scales across 3-dimensional volumes. Current optical approaches lack the spatial coverage and resolution necessary to measure and manipulate naturally occurring patterns of large-scale, distributed dynamics within and across deep brain regions such as the striatum. We designed a new micro-fiber array and imaging approach capable of chronically measuring and optogenetically manipulating local dynamics across over 100 targeted locations simultaneously in head-fixed and freely moving mice. We developed a semi-automated micro-CT based strategy to precisely localize positions of each optical fiber. This highly-customizable approach enables investigation of multi-scale spatial and temporal patterns of cell-type and neurotransmitter specific signals over arbitrary 3-D volumes at a spatial resolution and coverage previously inaccessible. We applied this method to resolve rapid dopamine release dynamics across the striatum volume which revealed distinct, modality specific spatiotemporal patterns in response to salient sensory stimuli extending over millimeters of tissue. Targeted optogenetics through our fiber arrays enabled flexible control of neural signaling on multiple spatial scales, better matching endogenous signaling patterns, and spatial localization of behavioral function across large circuits.

## INTRODUCTION

Neural dynamics supporting perception, decision making, action, and learning vary across multiple spatial scales in the mammalian brain, within and across brain regions. An ongoing technical challenge in systems neuroscience is to measure and manipulate cell-type and neurotransmitter specific dynamics at scale in the context of behavior. Towards this goal, wide field microscopy has enabled millimeter scale functional imaging with genetically encoded indicators and targeted optogenetic manipulations across nearly the entire superficial cortex of rodents^1–7^. These studies have revealed large scale signaling patterns correlated with simple and complex features of behavior and perception that vary in time over millimeters of brain tissue. Optogenetic manipulations designed to match the scale of these patterns have causally linked large scale cross and within region signaling with behavior^5^. However, optical access to rapid neural dynamics in brain regions deeper than ∼1mm is limited by scattering and requires tissue penetration^8,9^.

GRIN lenses or optical cannulae are commonly used for cellular resolution imaging and manipulations from deep brain regions, but these are limited to only small (∼0.5-1mm) fields of view and require significant damage to overlying tissue^8,10^. Alternatively, optical fibers can be used to deliver and collect light from deep brain regions, permitting fluorescent measurements (fiber photometry) and optogenetic manipulations of population signals or neurotransmitter release near the tip of the fiber^11,12^. While fibers do not provide cellular resolution, their diameters are significantly smaller than GRIN lenses, presenting a possible utility for accessing larger scale population dynamics. Moreover, the spatial resolution of fiber photometry is well suited for measuring regional variations in neuromodulator release with fluorescent sensors.

Typical fiber photometry and optogenetic manipulation experiments use only one or two optical fibers targeting a single brain sub-region. While fiber photometry experiments can provide information about dynamics in a targeted region in behavior, they provide a limited view of how dynamics are coordinated in space across functionally heterogeneous, extended circuits, such as the striatum. Single or dual fiber optogenetics experiments, similarly, do not enhance or impair the natural spatial patterns of activity and thus may result in misleading findings, particularly in the case of null results. Without prior knowledge of where task-relevant dynamics are localized, fiber photometry measurements and optogenetic manipulations may be incompletely or mis-targeted. Another complication of interpreting traditional photometry and optogenetics experiments is that standard fibers transmit and collect light from a relatively large volume around the fiber tip, limiting spatial resolution and perhaps obscuring functional heterogeneities within the collection volume.

Recent advances in multi-fiber approaches have attempted to address the problem of spatial scale by increasing the number of implanted fibers to 12-48^13,14^. However, these techniques employ conventional large diameter fibers (100-200µm) and thus are not capable of high density measurements or targeted manipulations across single brain structures without causing significant tissue damage. Standard fibers collect fluorescence over relatively large volumes around the tip, limiting spatial resolution^15^. Additionally, existing implant designs involve rigid, linear arrays with fixed spacing, require specialized connectors, and do not permit flexible targeting of arbitrarily spaced regions^13^. Limitations in scalability and spatial resolution of existing techniques are particularly problematic for investigating large deep brain structures such as the striatum, which is known to exhibit heterogeneous signaling and control diverse behavioral functions on spatial scales ranging from 100s of microns to millimeters^16–20^. Small diameter fiber implants offer the possibility of denser recordings but must address targeting and signal detection challenges^21^. Here we report a scalable new optical fiber array approach using small diameter fibers capable of measuring and manipulating robust signals from multiple fluorescent sensors at over 100 targeted locations simultaneously over deep 3-dimensional volumes in behaving mice.

## RESULTS

### Microfiber array design

To enable precise, high density targeting of many locations throughout arbitrarily sized deep brain volumes, we designed an implant capable of holding small diameter (37µm or 50µm O.D.) optical fibers with custom spacing in 3-dimensions (Figure 1A). We successfully tested implants holding 30-103 fibers. The total tissue volume displaced by the fiber arrays is comparable to single or dual implants of conventional large diameter (200 or 400µm) optical fibers and less than standard (0.5-1.5mm) GRIN lenses. We primarily designed implants to cover locations across the striatum volume, but arrays can be flexibly fabricated to cover any combination of desired brain regions. The smaller diameter fibers sample fluorescence from significantly smaller tissue volumes than conventional fibers, allowing for denser optical measurements at higher resolution^15^ (Supplemental Figure 1A). Fibers were separated by a minimum of 250µm axially and 220µm radially, ensuring no significant overlap between fiber collection fields^15^. Fibers were arranged through holes in a micro-3D printed, biocompatible plastic grid and proximal ends were glued within a polyimide tube then cut and polished to form a smooth imaging surface (Figure 1A-B). To enable optical measurements of cell-type and neurotransmitter specific signaling, mice were injected at multiple locations with AAVs to drive expression of genetically encoded fluorescent indicators (e.g. dLight1.3b^22^, GCaMP7f^23^, Ach3.0^24^, Supplementary Table 1).

**Figure 1:**
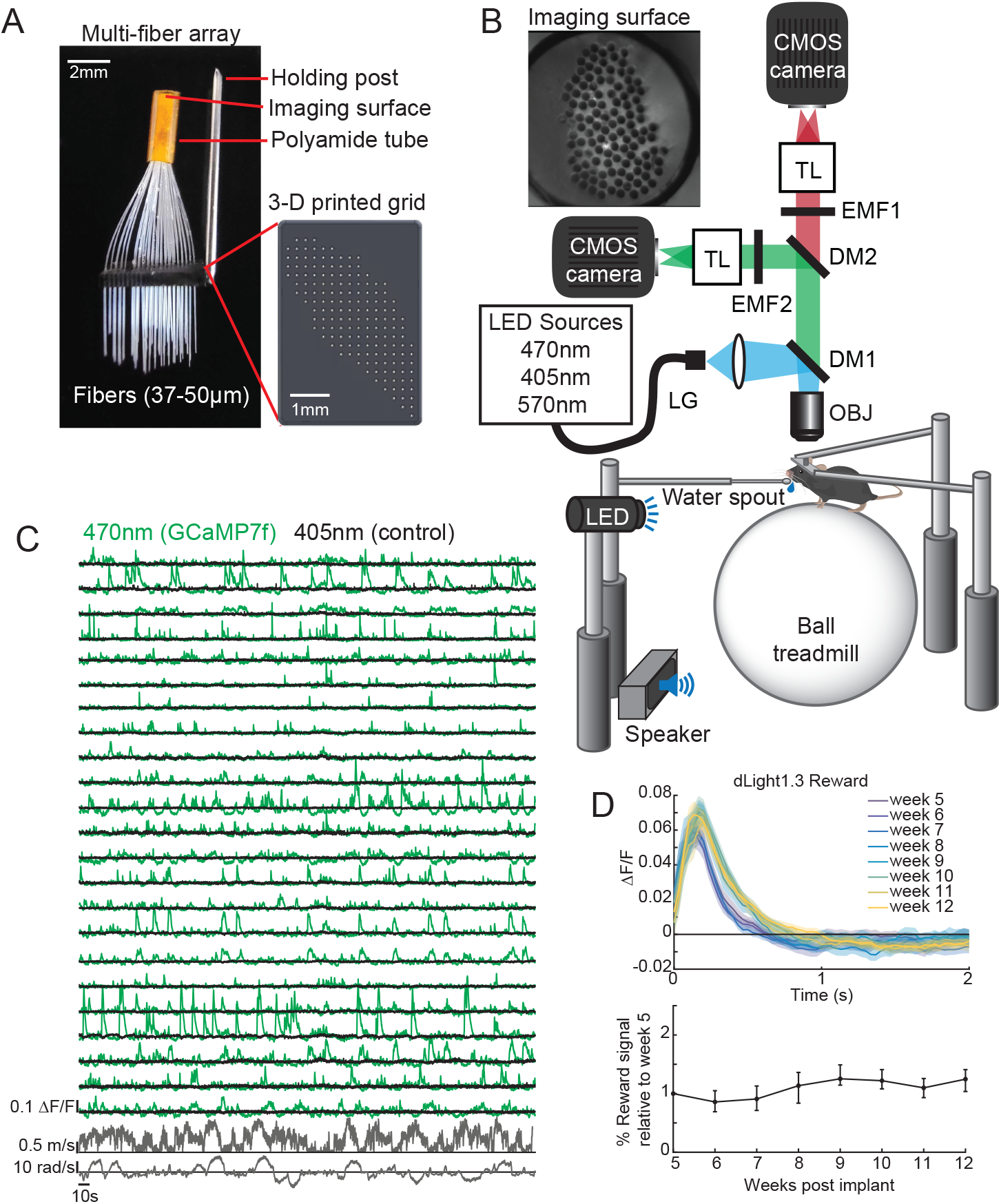
Multi-fiber arrays for large-scale optical measurements across deep brain volumes in behaving mice. **(A)** Multi-fiber array design. **(B)** Schematic of the microscope design for fiber bundle imaging (inset) in head-fixed behaving mice. TL, tube lens; EMF, emission filter; DM, dichroic mirror; OBJ, objective. See Methods for details on each component. **(C)** Changes in GCaMP7f fluorescence (ΔF/F) measured from 23 example fibers in a 93 fiber array implanted in the striatum of a D1-cre mouse. 470 nm excitation light (green trace) was alternated with 405nm light (black) at 18hz to provide a quasi-simultaneous isosbestic control. Significant transients are absent with 405 illumination. Bottom: angular and linear treadmill velocity. **(D)** Top: Mean changes in peak dLight1.3b fluorescence (DA release) from a single representative fiber aligned to unpredicted water reward consumption at different timepoints up to 12 weeks post-array implantation. Shaded regions, S.E.M. Bottom: Median peak reward ΔF/F across fibers (n = 44 fibers) in one mouse, normalized to initial responses on week 5. Upper and lower bars are the 75th and 25th percentiles respectively.

### Chronic imaging of large-scale calcium and neuromodulator signals in head-fixed and freely moving mice

Mice were outfitted with metal head-plates during array implantation to allow for fiber imaging via a table mounted microscope during head-fixed behaviors on a floating 2-D treadmill^25^ (Figure 1B). Illumination was provided by three high-power LEDs which could be alternated and synchronized with two sCMOS cameras to enable quasi-simultaneous imaging of red and green fluorophores (570nm and 470nm respectively, Supplemental Figure 2A) and an isosbestic control (405nm) for motion and hemodynamic artifacts. We initially validated the ability of our fiber arrays to reliably detect changes in cell-type specific Ca^2+^ signaling across the striatum using the genetically encoded sensor GCaMP7f expressed selectively in D1 spiny projection neurons (Figure 1C). Spontaneous Ca^2+^ transients during locomotion were observed across locations which were clearly separable from noise and highly heterogeneous across locations (Figure 1C). These transients reflected spatially weighted Ca^2+^ signals from neuron populations within a small volume around the fiber tips (Supplemental Figure 1A)^15,26–28^. Transients were not observed with illumination at the 405nm isosbestic point, indicating minimal contribution from hemodynamic or motion artifacts (Figure 1C). Robust, striatum-wide signals were also obtained with optical sensors for dopamine (DA) and acetylcholine (Ach, Supplemental Figure 2B) and for 37µm and 50µm diameter fibers (Supplemental Figure 1B). DA release at a single location to unpredicted water reward delivery was highly reliable and comparable in amplitude over at least 12 weeks following stable expression of dLight1.3b, confirming the utility of this approach for chronic measurements (Figure 1D).

To enable fiber array measurements in freely moving animals, we attached ‘miniscopes’^29,30^ above the fiber bundle implant and monitored fluorescence across fibers as animals explored a behavioral arena (Figure 2A). This revealed robust, high-SNR signals from GCaMP7f expressed in D1 spiny projection neurons that varied dynamically during locomotion (Figure 2B). Activity was highly heterogeneous, indicating complex activity dynamics across the volume of striatum. We also used miniscope imaging to monitor striatal DA release with dLight1.3b as mice foraged for food rewards. Interestingly, this revealed large transients that appeared to be reliably associated with the initial approach toward food rewards prior to consumption (Figure 2C).

**Figure 2:**
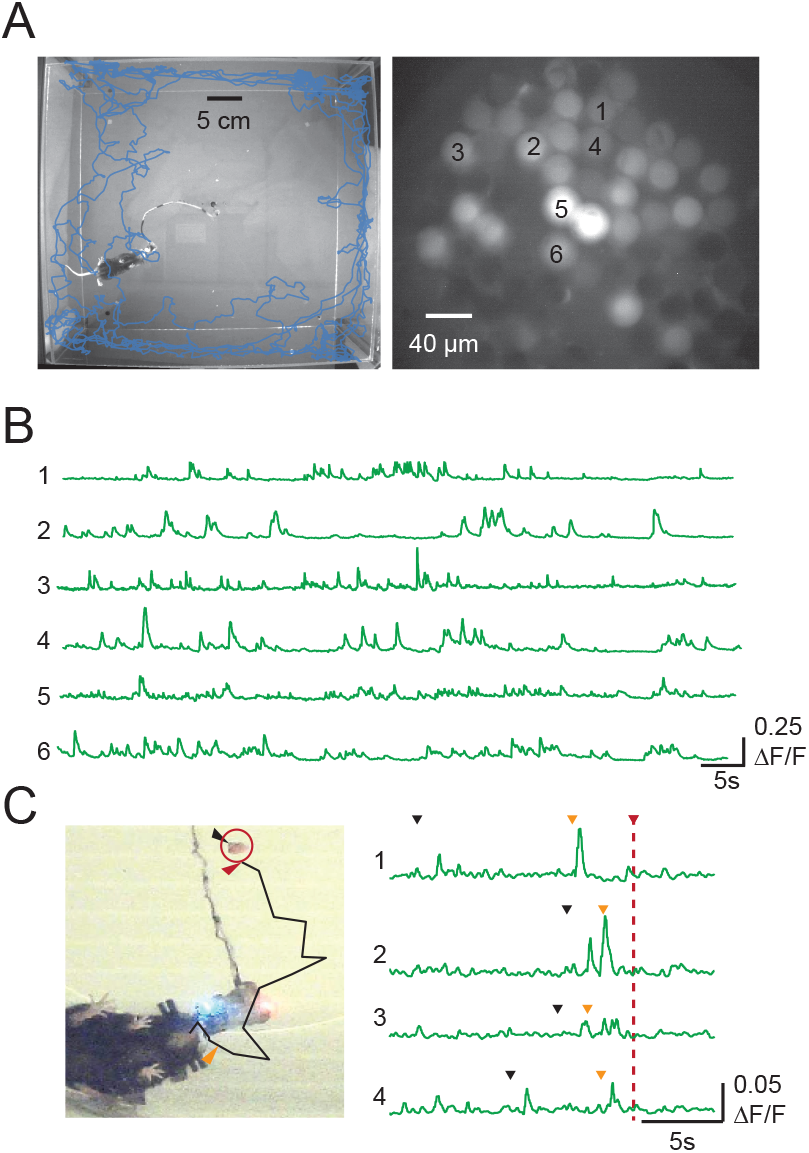
Miniscope imaging of multi-fiber arrays in freely-moving mice. **(A)** Locomotor trajectory of a miniscope-implanted mouse (left) and view of the fibers imaged with the miniscope (right). **(B)** Example GCaMP7f fluorescence traces (ΔF/F) from 6 fibers with robust signal in a D1-cre mouse. **(C)** Path of a mouse (left) converging on a food reward (red circle) during dLight1.3b (DA) imaging (right), along with example ΔF/F traces from one fiber over 4 food deliveries aligned to onset of consumption (red arrows). Black arrows indicate when the food was dropped into the arena; orange arrows indicate the estimated beginning of approach.

### Micro-CT based method for precise fiber localization and atlas alignment

As the fibers were densely spaced and smaller and more flexible than larger fibers, conventional histological techniques for fiber localization were not feasible. Therefore, we utilized a multi-step approach to first map each fiber on the array to the imaging surface pre-implantation (Figure 3A) then localize the tips of each fiber in the brain postmortem (Figure 3B-E). For postmortem localization, we used micro-computerized tomography (CT) scanning which allowed partially automated tracing of each fiber to the grid surface and precise identification of the fiber tip with approximately 10µm resolution (Figure 3B-C). Tissue visualization via CT was enabled by soaking the brains in a Lugol’s solution prior to scanning to enhance contrast^31^. CT images were scaled and aligned to a common coordinate framework^32^ (Figure 3B-D) to combine measures across mice (Figure 3B-E, see Methods).

**Figure 3:**
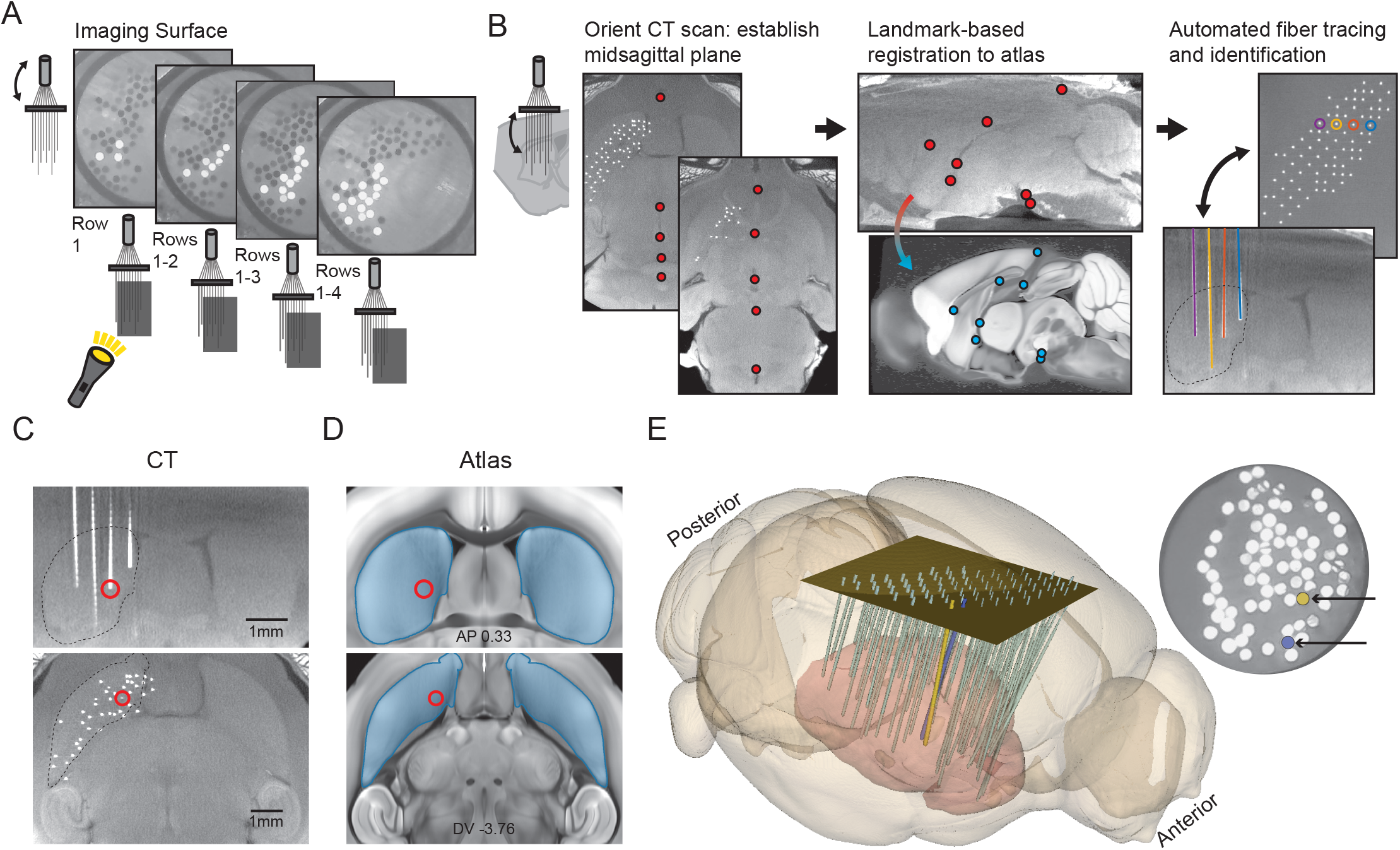
Calibration and post-mortem localization of micro-fiber arrays over deep brain volumes with computerized tomography. **(A)** Pre-implant calibration procedure for matching each fiber grid position with the corresponding location on the imaging surface. Example photos show the imaging surface as light is shone through subsequently uncovered rows at the distal implanted ends. This process was then repeated for columns. Images of illuminated fibers were used to match the fibers’ position on the grid surface (i.e., row and column location) with the imaging surface. **(B)** Schematic of the post-mortem localization procedure. Left: Micro-CT scans in the horizontal plane in two different example axial locations with manually placed markers (red dots) to identify the position of the midsagittal plane for proper orientation. Mid: Micro-CT scan in the horizontal plane with example manually placed markers for identified anatomical markers to enable registration to the Allen Mouse Brain Common Coordinate Framework atlas^32^ (bottom). Right: Micro-CT scans in the horizontal plane of the grid surface (top) and coronal plane (bottom) showing the tracing of 4 optical fibers. **(C)** Fiber tip localization. Micro-CT scan images in the coronal (top) and horizontal (bottom) planes from a mouse implanted with an 83-fiber array of 50µm fibers targeting the striatum. Red circles indicate the location of one fiber tip in both planes. **(D)** Coronal (top) and horizontal (bottom) plane atlas images corresponding to the locations of the registered CT images in c. Circles indicate the localized position of the highlighted fiber in **C**. **(E)** 3-D image showing atlas-aligned reconstructions based on CT images of every fiber in a 74-fiber array. Colored fibers (blue and yellow) crossed in the brain but were still unambiguously traceable back to the grid surface, and thus also the imaging surface.

### Spatial topography of modality specific dopamine release magnitudes over large-scale 3-dimensional volumes

To demonstrate the utility of our fiber array method to resolve large-scale spatial and temporal patterns of neural signaling, we measured striatum-wide DA release to salient stimuli using the fluorescent DA sensor dLight1.3b^22^ (Figure 4A). Midbrain DA neurons exhibit short latency responses to salient visual and auditory stimuli, which may drive learning, spatial attention, or orienting responses^33–35^. Fiber photometry studies have established that salient, high intensity stimuli evoke short latency increases in the posterior striatum and decreases in the anterior ventral^36^. However, standard approaches have not permitted investigation of sensory-evoked DA release at striatum-wide scales. Mice were presented with either visual (blue LED) or auditory (12kHz tone) stimuli at random intervals (4-40s) (Figure 2B). Surprisingly, the tone and light stimuli evoked distinct, modality specific, spatiotemporal patterns of DA release across striatum locations (Figure 4B-H). The light evoked transient increases in DA release across many locations with an amplitude gradient extending from the anterior striatum into the posterior tail, where increases were less prevalent and smaller amplitude (Figure 4E; Supplementary Figure 3). In contrast, the tone evoked the strongest increases in DA release at sites in the posterior tail, while anterior to the tail, the tone evoked primarily decreases in DA release (Figure 4D,F; Supplementary Figure 3) Decreases were slightly more prevalent in the anterior lateral and ventral striatum, with many sites exhibiting both increases to the light and decreases to the tone (Figure 4F). These distinct release patterns were consistent within and across mice (Supplemental Figure 3). Locomotor changes to stimuli were variable, bi-directional, and differed across mice and individual stimulus presentations, while DA release patterns to light and tone were consistent, indicating that modality-specific patterns were stimulus, not movement driven (Supplemental Figures 3, 4). Large fluorescence changes and spatial patterns were not observed with 405nm illumination, ruling out a significant impact of hemodynamic changes on the spatial DA release patterns (Supplemental Figure 5). These data indicate that representations of stimulus saliency in striatal DA release are spatially organized by stimulus modality and polarity across striatum regions.

**Figure 4:**
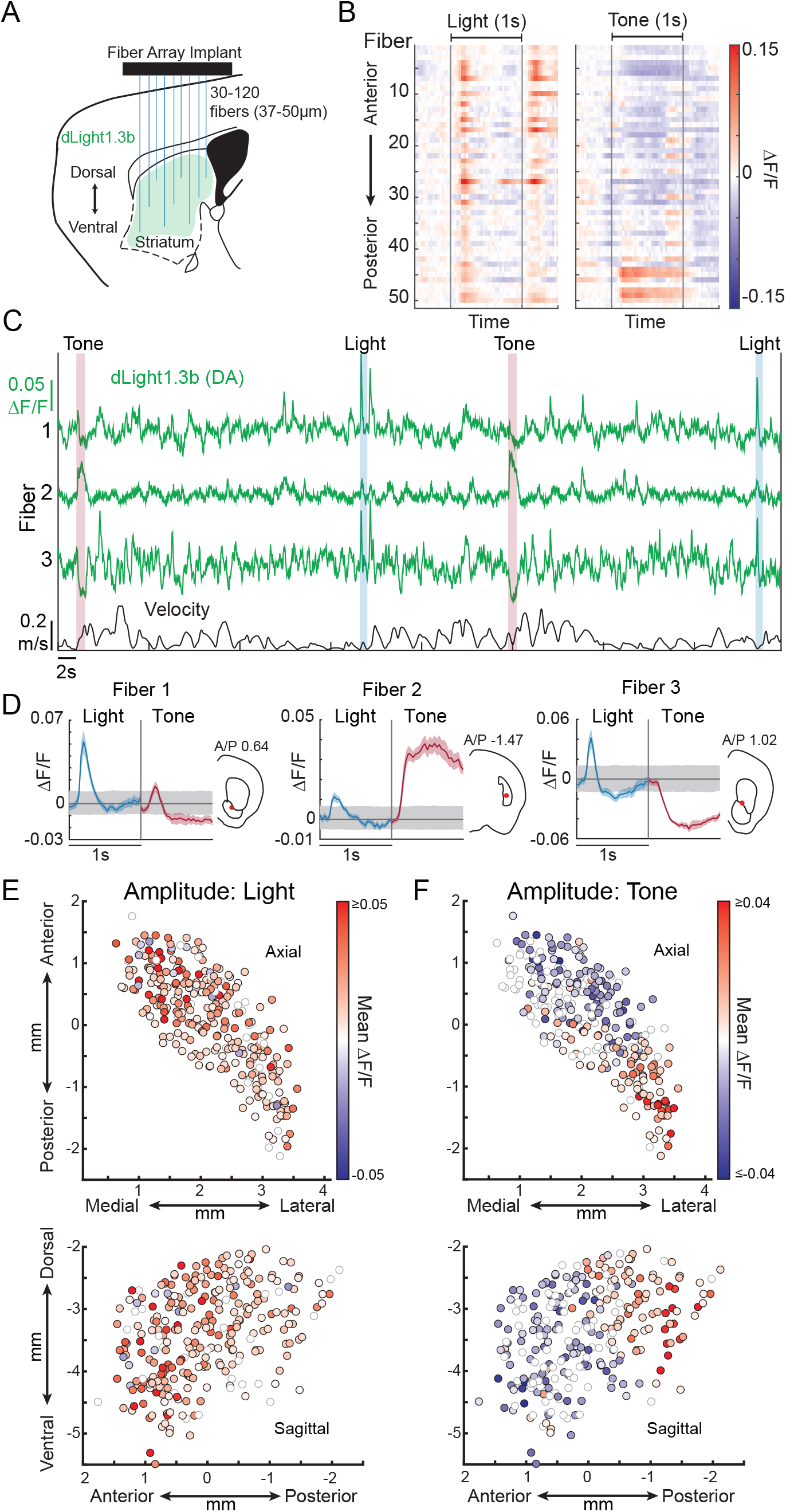
Modality specific patterns of striatum-wide dopamine release amplitude to salient stimuli. **(A)** Implant schematic for striatum-wide measurements of DA release with dLight1.3b. **(B)** ΔF/F (dLight1.3b) across all fibers (rows organized anterior to posterior from top to bottom) for single representative light (left) and tone (right) presentations. **(C)** ΔF/F traces from 3 example fibers from a single mouse (out of 280 fibers from 7 mice) during multiple presentations of light and tone stimuli. **(D)** Triggered averages of ΔF/F across all stimulus presentations (n = 18 each for light and tone) for the 3 fibers shown in 1c. Insets show brain locations of each fiber. Shaded regions, SEM. **(E)** Largest amplitude mean ΔF/F change across all light presentations for all fibers (n = 280) and mice (n = 7, 4F + 3M) in a 1-second window after stimulus onset. The color of each circle represents the largest mean ΔF/F change for a single fiber in either the positive or negative direction. The location of each circle indicates its localized fiber position in the axial (top) and sagittal (bottom) planes. Open circles, non-significant increases or decreases relative to baseline (mean exceeding 99% confidence interval of the bootstrapped null distribution for 3 consecutive timepoints, p<1.0×10^-6^). **(F)** Same as **E**, for tone presentations.

### Modality specific gradients of dopamine release timing across the striatum

The simultaneous parallel recordings and high spatial resolution afforded by our multi-fiber arrays enable investigation of how distributed neural signals are coordinated in time across neural space. A previous study has reported wave-like dynamics of reward-related striatal DA release in 2-dimensions in the dorsal striatum^37^. We assessed whether spatially organized temporal gradients were present for salient stimulus-evoked DA release across the 3-dimensional striatum volume. Response latencies (from stimulus onset to the half-max or half-min amplitude) were shorter, on average, for DA increases to light, and shorter for DA increases than for decreases to the tone but varied considerably (Figure 5A,B, light increase mean latency 221.6ms, SD 42.5; tone increase mean latency 305.4ms, SD 100.0; tone mean decrease latency 428.3ms, SD 197.1). We examined whether temporal structure was present across simultaneously recorded locations by comparing the trial-to-trial latencies of the DA responses to salient stimulus presentations. DA release latencies for light presentations followed a spatial gradient along the anterior-posterior axis, with relatively faster latencies to increase in the anterior striatum than in the posterior striatum, consistent with wave-like dynamics (Figure 5C, E, G). DA release latencies to tone presentations followed a different spatial gradient with faster latencies in the posterior striatum relative to the anterior striatum (Figure 5D, F, H). The posterior-anterior spatial gradient in the tone release latencies could be attributed to the decreases in the anterior striatum occurring at a longer latency than increases in the posterior tail (Figure 5B, Supplemental Figure 6). Subtle spatial structure along the M/L and D/V axes was observed for tone DA increase and decrease latencies in the anterior striatum and tail respectively when considered independently (Supplementary Figure 6). Spatial gradients were also present at the level of individual mice (Supplemental Figure 7). These data suggest that striatal DA release to salient stimuli propagates through the striatum with wave-like dynamics in modality-specific spatial gradients.

**Figure 5:**
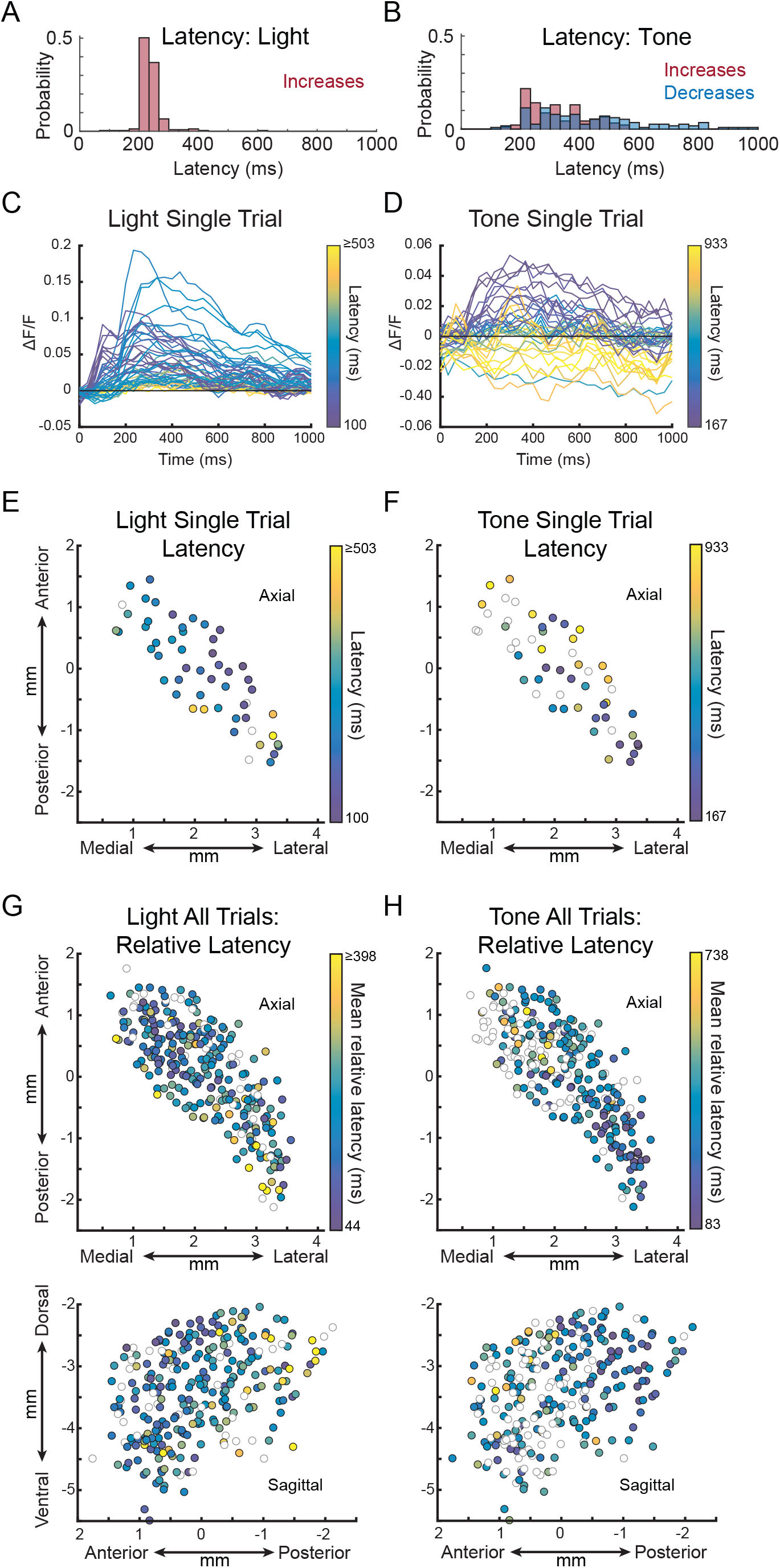
Temporal organization of modality specific striatum-wide dopamine release to salient stimuli. **(A)** Histogram of latencies to the half-max amplitude of the triggered average response to the light. **(B)** Histogram of latencies to the half-max amplitude of increases (red) or half-min amplitude of decreases (blue) of the triggered average response to the tone. **(C)** ΔF/F for a single light presentation across all fibers (n = 49/53) showing a significant response (single trial activity and mean triggered average exceeding 99% confidence interval of the bootstrapped null distribution for 3 consecutive timepoints, p<1.0×10^-6^). Traces for each fiber are colored according to the latency from stimulus onset to the half-max amplitude. **(D)** Same as **A**, but for a single tone presentation (n = 38/53 fibers). Latency was calculated from the stimulus onset to the half-max or half-min amplitude for fibers that increased or decreased respectively. **(E)** Spatial map in the axial plane of the half-max latencies from **C**. **(F)** Spatial map of the half-max or half-min latencies from **D**, as in **E**. **(G)** Spatial map of the mean relative half-max latencies of DA release for each fiber normalized to the minimum, half-max latency for each trial across all light presentations for all fibers (n = 280) and mice (n = 7, 4F + 3M, same mice as Figure 4). The color of each circle represents the mean relative latency for a single fiber. Open circles in spatial maps are non-significant increases or decreases relative to baseline (mean triggered average exceeding 99% confidence interval of the bootstrapped null distribution for 3 consecutive timepoints, p<1.0×10^-6^). **(H)** Same as **G**, for tone presentations. For fibers with significant decreases, the half-max latency to the minimum is shown (see Supplementary Fig 6).

### Targeted light delivery to individual micro-fibers for localized optogenetic manipulations

The DA release measurements above illustrate that bi-directional neuromodulatory signals that may modulate movement, perception, and learning are heterogeneously expressed across arbitrarily shaped 3-dimensional volumes in the striatum (Figures 4, 5). Precisely assessing the impact of large scale neuromodulator release on behavior requires targeted manipulations that match the spatial and temporal properties of the endogenous release, which is not possible with standard optogenetic manipulations with large diameter optical fibers. To enable flexible and targeted optogenetic manipulations of 3-D volumes, we incorporated a programmable digital mirror device (DMD, Mightex Polygon) into our microscope to target laser light (465 nm) into individual optical fibers (Figure 6A, B, D). We tested the ability of this system to control local striatum dopamine release by expressing the excitatory opsin channelrhodopsin-2 (ChR2) in midbrain DA neurons and the red DA sensor rDAm3.0^38^ across the striatum (Figure 6A). A fiber array was then implanted to target locations across the striatum volume. We confirmed that circular light spots could be targeted to individual optical fibers with no leak into neighboring fibers (Figure 6D). Light spot power at the objective (∼750 μW) was calibrated to achieve a power density at the fiber tip of 175 mW/mm^2^, which is estimated to activate ChR2-expressing axons in a volume extending approximately 150 microns axially from the bottom of the fiber tip (Figure 6C). This activation volume is significantly (>10x) smaller than the volume activated by the same power density through a standard 200μm, 0.37NA fiber (Figure 6C). A 570 nm LED was continuously projected onto the fiber bundle surface to excite rDA. The low power (1.6-2 mW/mm^2^ at the tip) of the 570nm excitation light and very low photoactivation of ChR2 at this wavelength ensured no stimulation during imaging.

**Figure 6:**
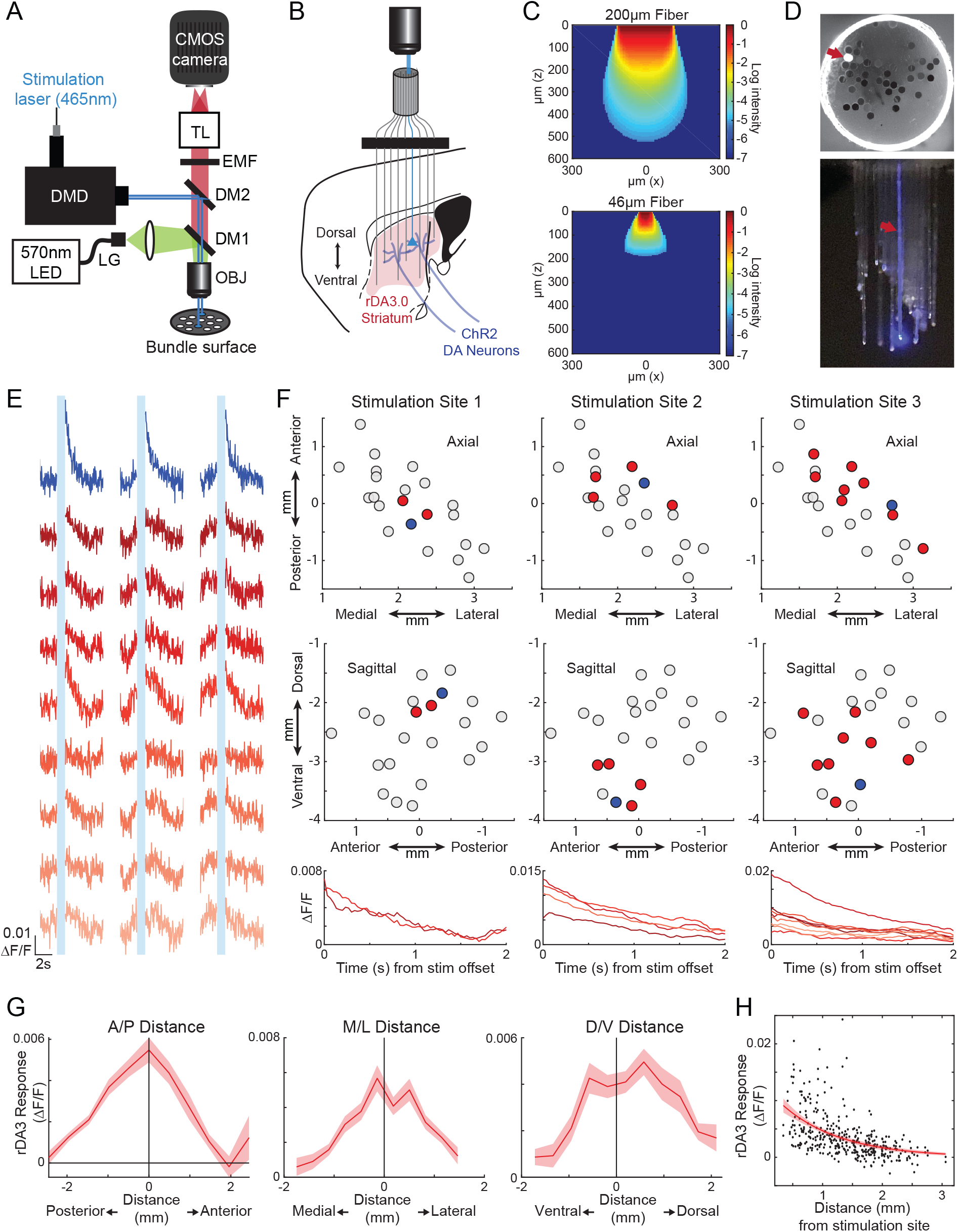
Light targeting to individual fibers permits localized optogenetic manipulations of striatal dopamine release. **(A)** Schematic of the microscope design and light path for simultaneous single site stimulation and imaging. TL, tube lens. EMF, emission filter, DM, dichroic mirror, OBJ, objective, LG, liquid light guide, DMD, digital mirror device (see Methods for details). **(B)** Schematic of injection and implant strategy. rDA3.0 was expressed throughout the striatum, while ChR2 was expressed in DA neuron axons. Fiber array implant targeted multiple sites throughout the striatum. Laser light was targeted to a single fiber to stimulate local DA axons. **(C)** Relative excitation light (465nm) intensity modeled as a function of distance from the optical fiber tip for a standard 200μm diameter, 0.37NA fiber (top) and the 50μm diameter fibers used in our arrays^45^ (bottom). Intensity values are shown only for locations estimated to achieve a power density sufficient to activate ChR2 (1mW/mm^2^). **(D)** Top: LED-illuminated fiber bundle surface with blue laser light targeted to a single fiber (red arrow). Bottom: distal, implanted ends of a fiber array. A single fiber is illuminated (red arrow), illustrating restricted targeting. **(E)** ΔF/F traces across fibers for three example stimulation trials at a single site (blue trace, top) and all other recording sites with a significant response to stimulation (bootstrap test, p<1.0×10^-6^), ordered from top to bottom by their total distance from the stimulation site (shaded from dark to light red). Vertical light blue bars indicate laser periods (4 ms pulses at 30 Hz for 1 second). **(F)** Spatial maps (axial, top; sagittal, middle) of significant changes in DA release (ΔF/F) for three example stimulation sites. Each circle is a single recording site, with the stimulation site in blue, sites with a significant evoked response (bootstrap test, p<1.0×10^-6^) in red, and sites lacking a significant response in gray. Bottom: mean ΔF/F triggered on stimulation offset at all of the sites with a significant response (left, n = 7 trials, middle, n = 9 trials, right, n = 9 trials). **(G)** Mean ΔF/F to single-site stimulation binned by distance from the illuminated fiber across 3 anatomical axes. Error bars S.E.M (n = 378; 21 recording sites per stimulation, 18 stimulation sites). **(H)** Mean ΔF/F for each non-stimulated fiber vs distance from the stimulated fiber for all stimulations. The red line shows the exponential model fit. Error bars 95% prediction confidence interval.

### Local control of striatal dopamine release through micro-fiber arrays

To measure the spread of evoked DA release in response to terminal stimulation through individual fibers, we delivered light bursts (1s duration, 30Hz, 4ms pulse width) to individual fibers while measuring DA release across neighboring locations. Large increases in fluorescence were observed at the site of stimulation, but these signals were partly contaminated by photoswitching of the rDAm3.0 sensor, as we observed smaller but significant increases in non-opsin expressing mice. Photoswitching in controls, but not opsin expressing mice, was entirely confined to the stimulated site (0/30 non-stimulated sites for 3 single-site stimulations, bootstrap test vs null distribution, see Methods), so we focused our analysis on the spread of stimulated release to non-stimulated fibers. Significant transient increases in stimulation-evoked DA release were detected at sites neighboring the targeted fiber which dropped off sharply with distance in all 3 dimensions (Figure 6G,H,J). Given the localized activation volume through our fibers (Figure 6C) relative to our fiber spacing (>350 micron separation), this DA release spread was not likely due to direct light propagation to neighboring fibers but to the highly branched morphology of DA axons^39^. Supporting this idea, activation dorsal to the stimulation location dropped off less with distance than ventral (Figure 6G, D/V Distance, right), which would not be expected based on the excitation profile (Figure 6C) but is consistent with the dorsally extending axon morphology of branching DA axons^39^. Thus, our fiber array method provides semi-localized control of neuromodulator release in the striatum enabling combinatorial stimulation to mimic the spatial and temporal spread of natural release to events such as salient stimulus presentation (Fig 4).

### Targeted optogenetic manipulations can localize region-specific behavioral effects in the striatum at multiple scales

Many large brain regions such as the striatum and hippocampus are composed of functionally distinct circuits and cell-types which provide parallel control over distinct aspects of behavior. In the striatum, for example, regionally restricted activation of D1 expressing spiny-projection neurons (dSPNs) can drive execution of specific actions^18^. Closely spaced circuits (<1mm separation), either within or across regions, cannot be independently manipulated by existing multi-fiber optogenetic approaches because of the broad light spread and the tissue damage caused by large diameter optical fibers (Figure 6C). To test whether we could apply targeted optogenetic stimulations to localize regions which control a distinct behavior, we stimulated ChR2-expressing dSPNs at individual locations targeted by our microfibers (Figure 7A). We focused on orofacial movements, as it is known that cortical orofacial projections terminate in distinct ventral striatal regions and that broad stimulation of D1-expressing neurons in those regions can modulate orofacial behavior^18,40,41^. Changes in orofacial movements were quantified by tracking the nose and points along the jaw in live video recordings using DeepLabCut^42^ (Figure 7B-C). To first confirm that we could drive region specific orofacial changes with broad unilateral stimulation, we coupled light into two groups of fibers targeting the ventral or dorsal striatum. We found that stimulating regions in the ventral, but not the dorsal striatum led to rapid increases in jaw movement, which were not present in the non-opsin expressing control mouse (2-way ANOVA interaction between region and optogenetic stimulation group p=0.011, two-sample t-test between optogenetic stimulation and control groups: p<0.01; p = 0.009, ventral; p = 0.394, dorsal, Figure 7D). These results confirm our ability to broadly localize effects of stimulation to specific striatum regions and are in agreement with the topography of orofacial inputs to the striatum and previous behavioral effects with larger diameter fibers^18,40^.

**Figure 7:**
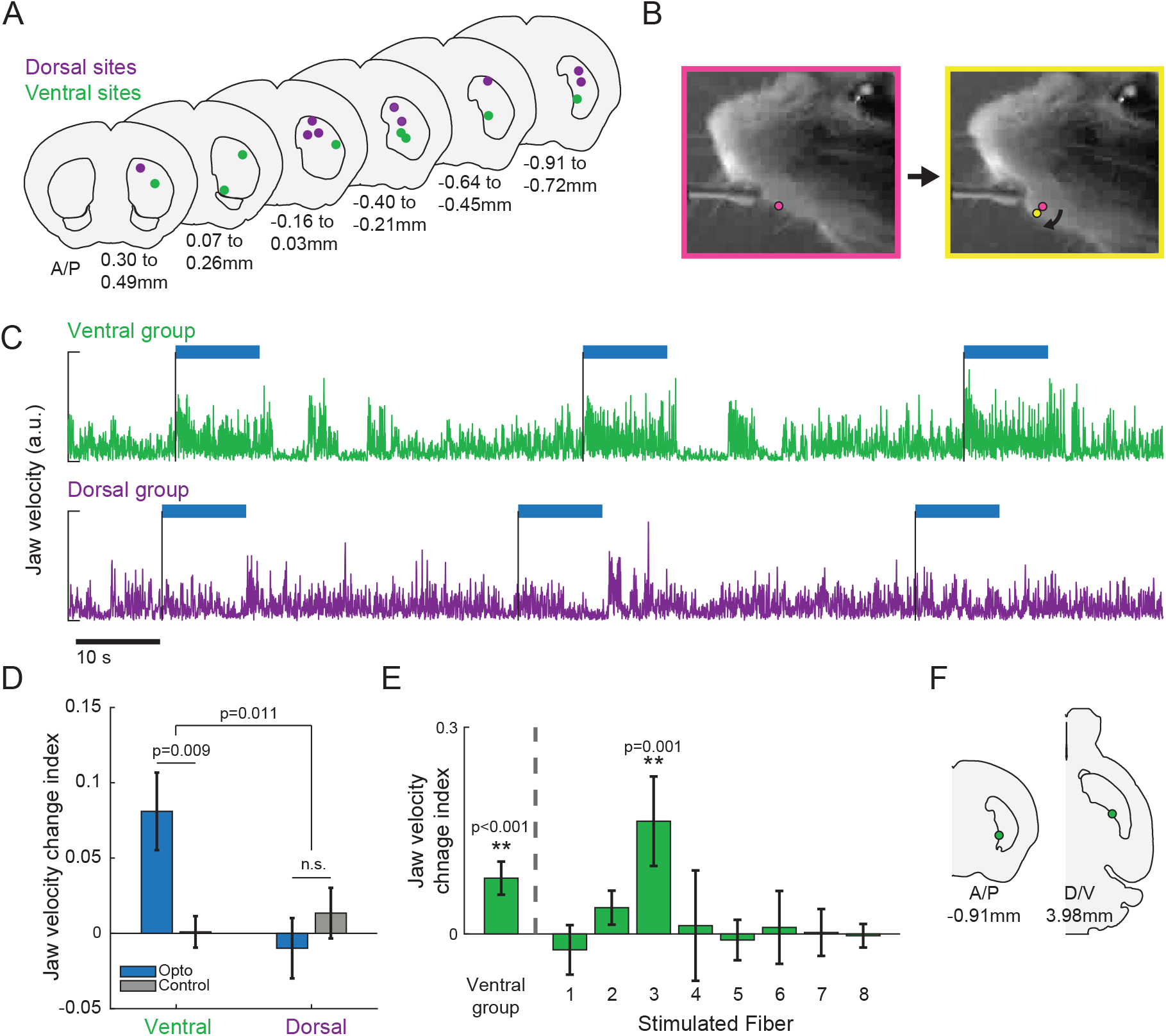
Localized effects of D1 spiny projection neuron stimulation on jaw movements. **(A)** Coronal cross-sections showing groups of stimulation sites targeted to the dorsal striatum (purple, n = 9) and ventral striatum (green, n = 8) in a mouse expressing ChR2 in D1 SPNs. **(B)** Two example live video frames separated by ∼66.66 ms, showing a change in jaw position during stimulation at all ventral sites in A. Pink and yellow markers indicate the front lower jaw marker, labeled automatically using DeepLabCut. **(C)** Jaw velocity in pixels/second during stimulation (blue bar, 4 ms pulses at 30 Hz for 10 s) delivered to a group of ventral sites (green, top) or dorsal sites (purple, bottom). **(D)** Average jaw velocity change index (see Methods) during stimulation at a group of ventral (left) and dorsal (right) sites in a ChR2-expressing (blue), and control mouse (gray). Error bars, S.E.M (from left to right: n = 47 trials; n = 39 trials; n = 48 trials; n = 42 trials). **(E)** Average jaw velocity change index (as in D) for the stimulation of all the ventral targeted sites simultaneously (far left) and each ventral site individually (n = 9 trials per stimulation site). Only one site had a significant change index when stimulated individually (**p<0.01, bootstrap test vs null distribution). Error bars, S.E.M. **(F)** Location in the striatum of fiber 3 in E (left, coronal view; right, axial view).

We then tested whether we could more precisely localize stimulation to regions driving jaw movements by coupling light into only single ventral striatum targeted fibers at a time. We found that only one of the eight ventral fibers included in the group stimulation produced a significant effect on jaw movements (Figure 7E). Post-hoc CT scanning (Figure 7F) localized this fiber to the posterior medial striatum, a region receiving heavy orofacial projections^40^. Other ventral striatum fibers which did not elicit jaw movements were located more anteriorly and medially, in regions which do not receive strong orofacial inputs (Figure 7A, E). Future experiments can use this approach to further probe the roles of striatal sub-circuits in controlling distinct actions. These results demonstrate the capability to conduct large-scale surveys of behavioral function across targeted locations within deep tissue volumes extending within and across closely spaced circuits.

## DISCUSSION

We developed and applied a new micro-fiber array and localization strategy that enables targeted measurements and manipulations of cell-type and neurotransmitter specific signaling across large, deep brain volumes during behavior and learning. The simple implant design and small diameter fibers allow for scaling well beyond existing multi-fiber photometry methods, while displacing comparable tissue. Implants can be easily modified to accommodate more (or less) fibers with any spacing by changing the configuration of the grid patterning (Figure 1A), without modifications to the microscope setup. Flexible scaling of fiber counts and spacing allows for investigation of within and cross-region network dynamics over many brain regions in parallel, and perhaps across brains of larger species such as rats or non-human primates.

Simultaneous, within animal measurements of widespread network dynamics can reveal the spatial scales over which behaviorally relevant dynamics are expressed. We applied the arrays to map the spread of distinct dopamine signals to salient auditory and visual cues across the striatum and describe a topography extending across millimeters in 3-dimensions (Figures 4, 5). Previous studies using single and dual fiber photometry have shown similar differences in dopamine release polarity between the anterior ventral and posterior striatum in response to salient tones but could not resolve how these signals extend across other striatal regions^36^. We also describe a new modality specific organization to the relative amplitudes and timing of stimulus-evoked DA release (Figures 4, 5). Simultaneous transmission of distinct, modality specific positive and negative DA release to downstream regions may differentially influence learning or immediate action to promote approach or avoidance of particular cues^43,44^. Understanding how neuromodulatory signals are balanced across regions, on immediate or long-term timescales, will likely be key to predicting their relative influence on downstream structures and ultimately their impact on behavior.

The smaller excitation and collection volume around each fiber tip^15,45^ (Figure 6C, Supplemental Figure 1) permits more localized interrogation of closely spaced circuits within or across deep structures. This feature is particularly important for large, heterogeneous structures such as the striatum or the hippocampus where circuit dynamics and behavioral function can vary over unknown scales ranging from hundreds of microns to millimeters^17,18,20,46–48^. The restricted collection volume and dense fiber spacing can also resolve spatiotemporal structure, such as ‘traveling waves’ or spatial synchrony across large volumes, which are not accessible by non-simultaneous, cross-group comparisons. Traveling waves across scales have been observed in multiple cell-types and systems and are hypothesized to play important roles in healthy and pathological circuit function ^11,49–51^. Indeed, waves of DA signaling have been described in two-dimensions across small regions of the dorsal striatum^37^. Our micro-fiber arrays provide an opportunity to resolve the propagation of waves accessible by fluorescent sensors and time varying synchrony across large volumes in 3-dimensions. Simultaneous multi-site measurements were applied to resolve 10s of ms timescale temporal offsets in modality-specific stimulus-evoked DA release across the striatum volume (Figures 4, 5). These results indicate wave-like patterns of bi-directional DA release which may reflect distinct signals that may differentially regulate learned and intrinsic responses to salient stimuli in downstream regions.

Drawing links between naturally occurring dynamics with behavior requires targeted circuit manipulations that accurately match the spatiotemporal characteristics of the dynamics in particular behavioral contexts. Optogenetics provides rapid, bi-directional control of specific cell-types within a region defined by the spatial profile of tissue illumination. Targeted light delivery to deep brain regions is typically achieved by 1-2, 100-200μm diameter optical fibers which activate opsins expressed in a tissue volume extending several hundred microns axially from the fiber tip^45,52,53^ (Figure 6C). Thus, depending on the geometry of underlying circuit dynamics, optogenetic experiments likely manipulate a volume which is either too small (in the case of widespread or distributed dynamics) or too large (in the case of sparse, localized dynamics). Moreover, most manipulations are conducted without knowledge of the spatial and temporal properties of the natural, underlying signals during the behavior of interest.

Our DA release measurements (Figures 4, 5) underscore the importance of considering the true spatiotemporal dynamics of signaling; Spatial profiles of modality selective, stimulus evoked DA release extended across 100s of microns to millimeters of space in 3-dimensions (Figure 4), and standard optogenetic manipulations of DA terminals or midbrain cell-bodies could not differentially probe the impact of these distinct, widespread signals on behavior without significant over or under targeting. We have demonstrated, using DMD guided light targeting of restricted locations through our fiber arrays, the ability to more precisely control the spatial spread and localization of optogenetic manipulations (Figures 6, 7). Importantly, we are able to combine manipulations with large-scale simultaneous measurements in the same animal to better match the spatial scale, magnitude, and locations of light delivery with natural dynamics (Figure 6). Future applications of this technique could utilize combinatorial illumination of multiple locations simultaneously to test the causal roles of distributed neural signals on behavior. Inversely, serial targeting of single, localized regions independently, within animal, could enable more precise localization of the critical circuit nodes that control specific behaviors (e.g. Figure 7).

## METHODS

### Multi-fiber array fabrication and calibration

Multi-fiber arrays were fabricated in-house to enable large scale measurements across deep brain volumes. Fibers (Fiber Optics Tech) had an outer diameter of either 37µm (34µm core, 3µm cladding) or 50µm (46µm core, 4µm cladding) with an 0.66NA. Bare fibers were cut into pieces (∼3cm) then mounted under a microscope into 55-60µm diameter holes in a custom 3D printed grid (Figure 1A, 3mm W x 5mm L, Boston Micro Fabrication), measured under a dissection microscope to target a particular depth beneath the grid, and secured in place with UV glue (Norland Optical Adhesive 61). Each array contained between 30 and 103 fibers separated by a minimum of 220µm radially and 250µm axially. Separation was calculated to achieve maximal coverage of the striatum volume with no overlap in the collection fields of individual fibers^15^. Distal ends were then glued inside an ∼1cm section of polyimide tube (0.8-1.3mm diameter, MicroLumen) then cut with a fresh razorblade. The bundled fibers inside the tube were then polished on fine grained polishing paper (ThorLabs,polished first with 6 micron, followed by 3 micron) to create a smooth, uniform fiber bundle surface for imaging. A larger diameter post was mounted on one side of the plastic grid to facilitate holding during implantation.

Prior to implantation, a calibration procedure was performed on the fabricated fiber arrays to match each fiber on the implanted side to a location on the bundle surface. To do this, distal (implanted) ends were illuminated with a flashlight, and transmitted light was imaged on the bundle surface. A sequence of illuminations was performed with different rows and columns covered, allowing each fiber on the imaging surface to be mapped to a specific row and column of the grid and distal fiber ends (Figure 3A).

### Stereotaxic viral injections and array implantation

C57BL6 (Figures 1-5, Supplemental Figures 3-7), DAT-cre (Figure 6, Supplemental Figure 2, Jackson Labs, strain #006660), or Drd1-cre (Figures 1, 2, 7, Jackson labs, strain #030329) ages 11 to 24 weeks were injected with AAVs to express genetically encoded proteins for optical measurements and manipulations (see Supplementary Table 1). Mice were anesthetized under isoflurane (1-3%) and a large craniotomy was performed with a surgical drill (Avtec Dental No. RMWT) to expose the cortical surface above the striatum. For striatum expression, virus was injected through a pulled glass pipette (diameter 30-50µm) at 10-40 total locations (200-800nL at each location) chosen to maximize overlap with fiber positions.

Injections were targeted to the midbrain for the stimulation experiment (Figure 6) at 4 sites relative to bregma: AP = -3.05, ML = 0.6, DV = -4.6 and DV = -4.25; AP = -3.5, ML = 1.25, DV = -3.9 and DV = -4.5. Injections to target midbrain DA neurons for the dual-wavelength recordings (Supplemental Figure 2B) were performed 3.07mm caudal to the bregma at four sites: ML = 0.5mm, DV =-4.00 mm and -4.25 mm, ML = 1mm, DV = -4.125mm and ML = 1.5 mm, DV = -3.8 mm below the dura, 200-800nL at each site at a rate ∼ 100 nL/min.The fiber array was then mounted in the stereotaxic and slowly lowered into position until one side contacted the skull surface. A thin layer of Kwik-Sil (WPI) was applied to seal off the exposed edges of the craniotomy, and then Metabond (Parkell) was used to secure the plastic frame to the skull. After this initial layer of Metabond solidified, a metal headplate and ring^20,25^ (Atlas Tool and Die Works) were secured to the skull with Metabond and the surface covered with a layer of blackened Metabond (carbon powder, Sigma). Finally, a small cylindrical plastic protective cap cut to extend just above the fiber bundle end was secured around the bundle and covered on the inside surface with blackened Metabond. Behavioral habituation began 1 or more weeks after surgery, and neural data acquisition began 3-5 weeks after injection and implant surgery.

### Micro-CT scanning and fiber localization

After completion of imaging and behavioral experiments, mice were injected intraperitoneally with 400-500 mg/kg Euthasol (Covetrus Euthanasia III), and then perfused transcardially with 20mL 1% phosphate buffered saline (PBS), followed by 20mL 4% paraformaldehyde in 1% PBS. After perfusion and decapitation, the lower jaw and front of the skull were removed in order to allow diffusion of solution into the brain while still keeping the implant intact. The brain was soaked in the 4% paraformaldehyde solution for 24h, rinsed three times with 1% PBS, and then transferred to a diluted Lugol’s solution, to provide tissue contrast for computerized tomography (CT) scanning^31^. The Lugol solution was prepared by diluting 10mL 100% Lugol’s Solution (Carolina, 10% potassium iodide, 5% iodine) with 30mL deionized water, a dilution chosen to be approximately isotonic to biological tissues^54^. Samples were soaked in this diluted Lugol’s solution in a foil-wrapped 50mL conical centrifuge tube on an orbital shaker plate for 10-14 days. We have more recently found that using 4 oz specimen cups instead of the 50mL conical centrifuge tubes enables better diffusion of the Lugol’s solution, and adequate contrast can be achieved in four days. After soaking, the skulls were rinsed three times with 1% PBS, secured in a modified centrifuge tube, and imaged in a micro-CT scanner (Zeiss Xradia Versa 520, a core instrument of the Boston University Micro-CT and X-ray Microscopy Imaging Facility) with the following parameters: 140kV, 10W, HE1 filter, 0.4X objective, 2s exposure time, 1001 projections, 12-micron voxel size.

The CT was then registered to an atlas to bring individual mice into a common coordinate system, and then fibers were identified and mapped from the recording tip up to the grid. To register the CT to the atlas, the 3D image was first oriented coronally, using 50-150 experimenter-identified points along the midsagittal plane, and a rough estimation of the anterior-posterior tilt of the brain. Then, previously-established landmarks^55^, were manually identified and marked on the CT scan using the Name Landmarks and Register plugin in ImageJ. A reference set of these landmarks was established by taking the mean location of the landmarks independently identified by seven lab members on the Allen Mouse Brain Common Coordinate Framework 3D 10-micron reference atlas^32^. All subsequent steps were carried out in Matlab (Version 2020b) using a combination of existing Matlab functions and custom lab-written functions and GUIs. The landmarks, separated into lateralized landmarks and midsagittal landmarks, were then used to register the CT to the reference atlas in a 3-step process. First, the midsagittal landmarks, in combination with the 50-150 previously identified midsagittal points, were used to re-establish the midsagittal plane, and the medial-lateral dimension of the CT was rescaled to match the atlas using the distance between the lateralized landmarks and midsagittal plane. Next, a rigid registration based on the midsagittal landmarks established a coarse anterior-posterior and dorsal-ventral alignment. And finally, the anterior-posterior and dorsal-ventral alignment was refined using an affine registration based on the midsagittal landmarks. The registration was then verified and manually refined using a lab-developed GUI.

Fibers were then identified and localized in a series of semi-automated steps. Fibers are very bright on the CT scan (Figure 3B-C) so are easy to identify automatically based on intensity. First, the tops of the fibers were aligned and assigned to the grid (which was previously mapped to the imaging surface). A dorsal slice was chosen where the row and column organization of the fibers could be clearly visualized. The fiber cross sections were automatically detected using the MATLAB functions *adaptthresh* and *imbinarize* to binarize the image, and then *bwconncomp* with a connectivity of 4 to find connected components, i.e., each fiber. The fibers were then automatically assigned to the grid based on spacing and input grid design, and the assignment was manually checked and refined as necessary.

Next, on an axial max projection of the CT scan, a polygon was drawn around the fibers to restrict fiber identification to this area. The CT scan was binarized, using *adaptthresh* and *imbinarize*, as before, within the defined area on each axial slice. The resulting binarized volume was then segmented into 3D connected components using *bwconncomp*, with a connectivity of 26. Only components connected to one of the identified, grid-assigned, fiber tops were considered fibers. In some cases when fibers crossed, they were lumped together into the same 3D connected component, and these cases were easily identified by virtue of being connected to multiple fiber tops. These “multifibers” were separated using watershed segmentation (*watershed* function in Matlab) on sequential axial slices to separate the touching fibers. For each of the fiber tops in the “multifiber”, a line was established from the centers (found using the *regionprops* function in Matlab) of each of the fiber’s cross sections in the first 10 slices. Given the voxel size, and fiber spacing of at least 220 microns in either the medial-lateral or anterior-posterior directions, it is not possible for them to cross within these first 10 axial slices. Continuing to move ventrally, watershed-separated fiber cross sections were assigned to each “fiber line” based on best fit, and with each subsequently assigned section, the line was iteratively updated. The resulting separated “multifiber” was verified by the experimenter and refined as necessary. Once there was a 1:1 mapping between fiber tops and fiber bottoms, the fiber identification step was complete, and the recording location was assigned to the center of the most ventral cross-section slice of the fiber. Anatomical labels were automatically assigned to each fiber recording location using available segmentations of the Allen Mouse Brain Common Coordinate Framework 3D 10-micron reference atlas^32,56^. Assigned locations were then manually verified and anatomical labels were refined in borderline cases where necessary.

For approximate conversion of recording locations in 3D atlas space to a Bregma-centered coordinate system, we converted distance in voxels (isotropic 10 microns) to distance from Bregma. The medial-lateral coordinate of Bregma was defined as the midsagittal plane. The anterior-posterior coordinate of Bregma was taken from an open-source atlas that merged the Franklin-Paxinos atlas into the Allen Mouse Brain Common Coordinate Framework 3D atlas^56^. The dorsal-ventral coordinate of Bregma was estimated by taking the average of repeated identification of landmarks along the midsagittal line on the AP=0mm coronal slice on both the Allen Mouse Brain Common Coordinate Framework 3D atlas and the Franklin Paxinos atlas, and then from those points, estimating the 0 point on the Allen Mouse Brain atlas. We note that the Allen Mouse Brain Common Coordinate Framework 3D atlas does not provide Bregma coordinates due to methodological considerations^57^. When provided, our Bregma-centric coordinates are for purposes of approximate reference for comparison with other atlases.

### Multi-color fiber array imaging

Fiber bundle imaging for head-fixed experiments was performed with a custom microscope mounted on a 4’ x 8’ x 12’’ vibration isolation table (Newport, Figure 1B). Excitation light (470nm, 405nm, and 570nm) for fluorescent sensors was provided by three high-power Solis LEDs (Thor labs, No. SOLIS-470C, SOLIS-405C, SOLIS-570C) which were combined using a series of two dichroic filters (Chroma No. ZT532rdc and 405rdc). Light from the LEDs was filtered (Chroma No. ET405/10, ET555/25, ET473/24) and coupled into a liquid light guide (Newport No. 77632) with lenses (f = 60mm and 30mm, Thor labs No. LA1401-A and LA1805) and a collimating beam probe (Newport No. 76600). The liquid light guide was coupled into a filter cube on the microscope and excitation light was reflected into the back aperture of the microscope objective (10x, 0.3NA, Olympus No. UPLFLN10X2) by a dichroic beam-splitter (Chroma No. 59009bs). Light power measured at the focal plane of the objective was set to 65-90 mW which produced ∼1.6-2 mW/mm^2^ power at the fiber tips. Fluorescence from the fiber bundle was collected by the objective then passed through the dichroic beam-splitter used to direct the excitation light. A second dichroic (Chroma, No. 532rdc) reflected green and passed red fluorescence, and bandpass filters for red and green (Chroma, No. 570lp and 525/50m respectively) blocked residual excitation light and autofluorescence. A tube lens in each path (Thor labs, No TTL165-A) focused emission light onto the CMOS sensors of the cameras to form an image of the fiber bundle (Hamamatsu, Orca Fusion BT Gen III). The microscope was attached to a micromanipulator (Newport No. 9067-XYZ-R) to allow fine manual focusing and mounted on a rotatable arm extending over the head-fixation setup to allow for coarse positioning of the objective over the mouse.

Imaging data was acquired using HC Image Live (Hamamatsu). Single wavelength excitation and emission was performed with continuous, internally triggered imaging at 30Hz. For dual-wavelength excitation and emission, two LEDs were triggered by 5V digital TTL pulses which alternated at either 11Hz (30ms exposure) or 18Hz (20ms exposure). To synchronize each LED with the appropriate camera (e.g. 470nm LED excitation to green emission camera), the LED trigger pulses were sent in parallel (and decreased to 3.3V via a pulldown circuit) to the cameras to trigger exposure timing (20ms exposure). The timing and duration of digital pulses were controlled by custom MATLAB software through a programmable digital acquisition card (“NIDAQ”, National Instruments PCIe 6343). Voltage pulses were sent back from the cameras to the NIDAQ card after exposure of each frame to confirm proper camera triggering and to align imaging data with behavior data (see below).

### Optical setup and parameters for targeted optogenetic stimulations

To couple light into individual optical fibers in our array for targeted optogenetic manipulations, simultaneously with imaging (Figures 6, 7), we integrated a programmable digital mirror device (DMD, Mightex Polygon1000 Pattern Illuminator DSI-K3-L20) into the light path of our imaging microscope (Figure 6A). Excitation light was provided by a 3.2W, 465nm laser (Mightex, LSR-040-0465), which was coupled to the DMD with an optical fiber. Light from the DMD was coupled into the objective path by a dichroic (Chroma 570lpxr). 570nm excitation and emission filters (see ‘Multi-color fiber array imaging’ above and Figure 6A) enabled simultaneous imaging of red fluorophores during stimulation.

Control of light patterning and stimulation parameters was achieved with PolyScan2 control software (Mightex) and custom MATLAB functions. A calibration step was performed prior to stimulations to align the camera view with the PolyScan2 software, allowing us to design patterns of circular light (∼40μm diameter for each spot) to target individual fibers (Figure 6A, B, D). Transmission efficiency through 50μm diameter fibers in our arrays was calculated at approximately 39% based on comparisons between power at the objective and transmitted light through individual fibers. For stimulations of dopamine release (Figure 6), a light spot of ∼750 μW (measured at the objective) was used, resulting in an estimated power density of 175 mW/mm^2^ (0.29 mW total power) at each implanted fiber tip. For stimulations of D1 expressing neurons (Figure 7), a light spot of ∼600 μW at the objective was used, resulting in an estimated power density of 141 mW/mm^2^ (0.23 mW total power) at the fiber tip. We estimated relative excitation light intensity and excitation area as a function of distance from the fiber tip (Figure 6C) by applying a light scattering model developed by Yona et. al 2016^45^ with a scattering coefficient of 140 cm^-1^ approximated for the striatum based on Azimipour et al 2014^58^ and Al-Juboori et al. 2013^59^ and an activation threshold for ChR2 of 1 mW/mm^2^ ^60,61^.

Light pulse trains for stimulation (30Hz, 4ms pulse width, 1s or 5/10s durations for DA and D1 neuron stimulation respectively) were programmed in PolyScan2 and triggered with 5V digital pulses controlled via MATLAB and sent from a NIDAQ board (National Instruments PCIe 6343) to the Polygon. Stimulations were triggered manually every 30-60 seconds for 5-10 minutes at a time for all stimulation experiments.

### Freely moving imaging

For visualizing fibers in behaving mice (Figure 2), custom-built miniscopes^29,30^ were positioned above the fiber bundle, focused with an XYZ manipulator, and cemented in place with Metabond (Parkell) under brief isoflurane anesthesia. Mice were allowed to recover for 24-48 hrs after attachment, after which imaging was performed in a custom built acrylic arena as mice foraged for Froot Loops (Kellog). Both behavioral video and fiber signals were acquired at 30 Hz using custom Matlab acquisition software. Neural imaging data were acquired from DAQ and CMOS PCBs from the UCLA Miniscope project^62^. Illumination intensity was delivered by a 470 nm LED (Luxeon Rebel) at an intensity of TK-TK mW, controlled by a D-A interface (National Instruments) and BuckPuck LED controller (LEDdynamics, Inc). Imaging data were motion-corrected with NormCorre^63^, corrected for minor background light leak using spatial high-pass filtering, and converted to ΔF/F. Intensity timeseries for each fiber were extracted using manually selected circular ROIs. Locomotor activity was quantified using manual tracking in ImageJ.

### Head-fixed behavior

After grid-implant surgery, the mice recovered for at least one week before handling and behavioral habituation. For reward experiments, mice began water scheduling at this time, during which they received 0.8-1.5mL water daily, calibrated so they could maintain a body weight 85-90% of their free-water body weight, as described previously^20^. Mice were headfixed over a hollow styrofoam ball treadmill (Figure 1B, Smoothfoam, 8in diameter), supported by air in a 3D-printed plastic cradle, which allowed them to run in all directions^25^. To monitor the movement of the ball, two optical mouse sensors (Logitech G203 mice with hard plastic shells removed, mounted on posts) were mounted on posts, one at the back and one 90 degrees off to one side of the ball, with the sensors level with the ball’s “equator”. Pitch and yaw (y- and x-displacement, respectively) were read from the optical mouse sensor at the back, and roll (y-displacement) was read from the optical mouse sensor at the side. The optical mouse sensitivity was set to 400 dpi, polling rate 1kHz. Each optical mouse was connected to a Raspberry Pi (3B+), running a continuous multi-threaded python program to read in the continuous 1kHz change in x- and y-position (in dots), and output a proportional continuous voltage at 100Hz. The pixels-to-voltage conversion was set such that velocity magnitude of 3.5 m/s^64^ corresponded to the maximum output voltage of 3.3V. This velocity magnitude voltage was converted to an analog voltage via a digital-to-analog converter (DAC, MCP4725), and read in through an analog input pin on the NIDAQ board. The sign (direction) of the velocity was sent as a digital binary variable through a separate Raspberry Pi output pin and read in through a separate input pin on the NIDAQ board.

For unpredicted water reward delivery, a water spout mounted on a post delivered water rewards (9 μL, Figure 1B) at random time intervals (randomly drawn from a 5-30s uniform distribution) through a water spout and solenoid valve gated electronically. Licking was monitored by a capacitive touch circuit connected to the spout. Sound stimuli were presented via a USB speaker placed on the table approximately 20 cm in front of the ball (Figure 1B), and calibrated to deliver tones at varying intensities from 80-90 dB, as measured from the location of the mouse. The noise generated by the air supply under the ball treadmill was measured to be approximately 78-80 dB. Light stimuli were presented via a LED (Figure 1B, Thor labs, M470L3) mounted on a post level with the mouse, approximately 20 cm away from the ball, 45 degrees contralateral to the implanted side calibrated to deliver light at varying intensities from 1-27mW, as measured just in front of the LED. Salient stimuli were presented in randomized order and at varying intensities, 14-21 presentations of each modality, with intertrial interval randomly drawn from a uniform distribution of 4-40 seconds.

Data was input (licking, velocity, TTLs, etc) and output (to trigger stimulus delivery, reward delivery, LED, image acquisition, etc) at 2kHz by a custom Matlab program via the NIDAQ card. For synchronization of behavioral data with imaging data, TTLs were sent from the cameras to the NIDAQ card 500μs after the beginning of readout for each frame (Hamamatsu HC Live VSYNC). Behavioral data was then downsampled to match the sampling rate of the neural data by block averaging, i.e., behavioral data was averaged for each frame.

To record the movements of the jaw (Figure 7B), video recordings were taken using a Blackfly S USB3 camera, positioned to focus on a side view of the mouse’s face. The behavior cameras were triggered on the TTL output from the imaging cameras, as described above.

### Data analysis

Data were processed and analyzed using built-in and custom lab-written functions in Matlab (Version 2020b).

### Velocity and acceleration

Velocity signals from the optical mice were converted back to m/s for the pitch (forward/backward) and roll (sideways) directions, and to degrees/s for the yaw (rotational) velocity. Pitch and roll combined into a linear velocity measure 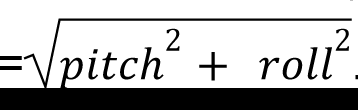. To get a measure of overall vigor, we used an “overall velocity” measure, for which we took the sum of the linear velocity and absolute value of the angular velocity (converted to m/s based on the ball circumference), low-pass filtered at 1.5Hz, and then smoothed with a 300ms smoothing window. Overall acceleration was calculated as the moment-to-moment difference of this velocity measure, low-pass filtered at 1.5Hz, and then smoothed with a 300ms smoothing window.

### Motion correction, ROI Selection, and calculation of ΔF/F fluorescence signal

Time series movies were motion-corrected using algorithms described previously^20^. Briefly, movies were then visually inspected to confirm image stability, and excluded from analysis in cases with excessive motion artifact. Mean fluorescence was extracted from circular regions of interest (ROIs) drawn for each fiber. ROI radius was approximately half that of the fibers (∼25-micron diameter ROIs for 50-micron fibers, and ∼19-micron for 37-micron fibers).

The same ROI map, adjusted to fit each movie, was used for the entire experiment, to ensure consistent ROI identification and mapping over multiple recordings. The mean extracted fluorescence traces were normalized to a baseline (8th percentile fluorescence over a 30s sliding window), to remove any slow frequency changes due to baseline drifts.

### Relationship of dLight1.3b fluorescence signals to salient stimulus presentations

To assess the relationship of fluorescence signals to the 1-second salient stimulus presentations while accounting for locomotion changes, peri-event (−1s to 2s around stimulus onset) fluorescence was modeled using a general linear model with overall velocity and acceleration predictors, and stimulus presentation event predictors convolved with a 13 degrees-of-freedom regression spline basis set with 1s duration, generated using Matlab’s *spcol* function. Velocity and acceleration predictors consisted of continuous overall velocity or acceleration at a lag estimated from cross-correlation analyses (+/-500ms lag) of fluorescence vs velocity, and fluorescence vs acceleration, during ITI periods. Velocity and acceleration were then entered as continuous predictors at the lag(s) corresponding to the most positive correlation coefficient (if significant, p<0.05) and the most negative correlation coefficient (if significant, p<0.05). For each fiber, then, there were 0-4 locomotion continuous predictors (0-2 for velocity, and 0-2 for acceleration), and 26 spline predictors (13 for each stimulus modality).

We therefore used Matlab’s *fitglm* to fit the following model to each fiber:

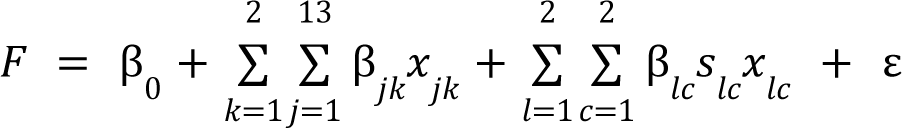

Here, *F* is the ΔF/F fluorescence signal from a single fiber, *k* = the stimulus modality (light or tone), *j* = the spline basis function, and *x_jk_* is therefore the *j*^th^ spline basis function convolved with a binary trace that is equal 1 at the times that the onset of stimulus *k* occurs, and 0 elsewhere. In terms of the locomotion continuous variables, *l* is the locomotion measure (velocity or acceleration), and c is the relevant cross correlation lags (the lag with the lowest negative correlation coefficient and the lag with the highest positive correlation coefficient); *s_lc_* is a binary variable indicating whether the correlation is significant (p<0.05) at that lag, and *x_lc_* is the corresponding locomotion continuous variable (velocity or acceleration) at the specified lag.

The resulting salient stimulus event spline predictors and coefficients were then reconstructed into an estimated event-triggered average. Significance of the event-related signal change was determined by whether this estimated event-triggered average exceeded the 99% confidence interval of a bootstrapped null distribution (10000 iterations) for 3 consecutive timepoints (p<1.0×10^-6^). To generate the bootstrapped null distribution, we fit the same model using randomly chosen salient stimulus onsets (same number of trials as actual events) instead of the actual event onsets, and reconstructed the estimated event-triggered average, repeated for 10000 iterations.

In Supplemental Figure 4, stimulus presentation trials were separated into trials followed by an acceleration and trials followed by a deceleration, determined by whether the maximum acceleration magnitude during the 1-s stimulus presentation was positive or negative, respectively.

The latency response was determined as the time elapsed between the beginning of the stimulus presentation when the signal surpassed the half max amplitude (half of the peak or dip amplitude, as applicable) either for the triggered average (Figure 4G-H) or on a trial-by-trial basis (Figure 5A-D). Trial-by-trial signals were reconstructed by regressing out the locomotion, as estimated by the GLM above. Trial-by-trial signals were considered significant if they met 2 criteria: first, the estimated event-triggered average was significant (Figure 4 and as described above), and second, the single-trial response also exceeded the 99% confidence interval of a bootstrapped null distribution (the same null distribution used for the mean) for 3 consecutive timepoints. If the signal crossed back and forth across the half max amplitude threshold multiple times before its peak, the crossing closest to the peak (i.e., the last crossing before the peak) was considered. The trial-by-trial relative latencies were calculated by subtracting the latency of the first fiber, such that the fiber with the fastest latency had a relative latency of 0.

### DA release to optogenetic stimulation

To determine the dopamine response to 1 s laser pulses delivered to individual fibers (Figure 6E), we first computed an event-triggered average of the ΔF/F trace for each recording location from 0-0.5 s from the offset of the laser. A significant response was determined by whether this event-triggered average exceeded the 99% confidence interval of a bootstrapped null distribution (10000 iterations) for 3 consecutive timepoints. The bootstrapped null distribution was determined by computing an event triggered average from n randomly selected timepoints, excluding the 1 s laser pulses, where n is the number of stimulation trials, repeated for 10000 iterations. The triggered average plots (Figure 6F) show the event triggered average from 0-2 s from the offset of the laser, smoothed using a moving average window of 0.3 seconds.

In order to determine the relationship between the stimulated response and distance from the stimulated location in each dimension (Figure 6G), we first computed the average ΔF/F for each recorded location and trial (inclusion criteria described below) within 0-0.5 seconds after the offset of the 1 s laser pulses. We then averaged these responses across trials for each stimulated fiber. The distance vector from the tip of the stimulated fiber to the tip of the recorded fiber in each dimension was computed by subtracting the position (determined from the CT scan as described above) of the stimulated fiber from the position of the recorded fiber. We then pooled all the responses for each stimulation (inclusion criteria described below), and binned them by distance into 12 evenly distributed bins for each dimension (bin widths, ML: 0.33 mm, AP: 0.49, DV: 0.38), and averaged the responses within each bin. Error bars show the standard error of the mean.

To determine the relationship between the stimulated response and the distance between the recording location and stimulation location (Figure 6H), we fit an exponential regression model using matlab’s *fit* function of the form:

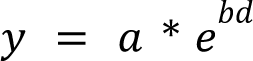

Where 𝑦 is the stimulated response, calculated as described above, and 𝑑 is the euclidean distance between the position of the recording location and the stimulation location (i.e., the length of the distance vector), calculated using matlab’s *norm* function:

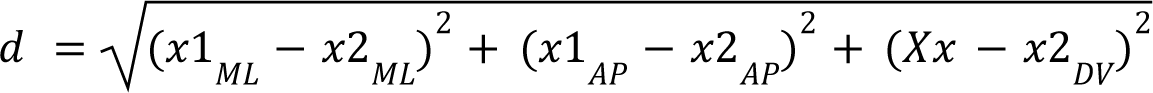

Where 𝑥1 is the position of the recorded fiber and 𝑥2 is the position of the stimulated fiber. Error bars show the 95% confidence interval of the predicted curve.

Inclusion criteria for the above analyses was three-fold. First, to be included in any of the analyses, the position of the fiber tip had to be within, or on the border of, the striatum, determined as described above (24 recording locations). Second, to confirm expression of the opsin, we only included stimulated locations for which 1 second laser pulses evoked a significant rDA3 response in at least one other location (18 out of 24 included locations). This was because a significant response at the stimulation site could be due to photoswitching of the rDA3. Third, to confirm expression of the sensor, we only included recording locations for which a significant response (defined above) was observed for at least one set of stimulations (22 out of 24 included locations).

### Behavioral response to optogenetic stimulation

In order to measure movements of the jaw in response to optogenetic stimulation of D1 neurons, a Python based toolbox called DeepLabCut^42,65^ was used. The side-view recordings of the mouse’s face (Figure 7B) were uploaded into Google Drive to be processed using DeepLabCut via Google Colab. Six points (spout, nose, upper jaw, front lower jaw, middle lower jaw, and back lower jaw) were manually labeled in random frames (selected using k-means algorithm) from 10 training videos across 3 different mice. The model was trained using the pretrained ResNet50 network for 300,000 iterations or until the loss plateaued. Low certainty labels were then manually corrected and merged with the previous training set to be trained again for another ∼300,000 iterations. The rest of our videos were labeled automatically using this final trained model. To quantify the jaw movement, we used the position of the front lower jaw as it appeared to be the most isolated marker for the jaw. The jaw velocity was calculated as the total displacement (in pixels) per frame:

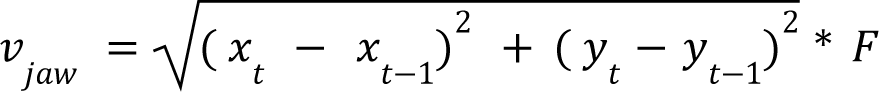

Where x and y are the coordinates of the front lower jaw marker at time t. We only included timepoints in which the likelihood of the DeepLabCut label was greater than 0.95 for at least 10 frames in a row. In frames with a likelihood of less than 0.95 the jaw was typically obscured by the mouse’s paw or tongue. The change in jaw movement due to stimulation was assessed via a jaw velocity change index:

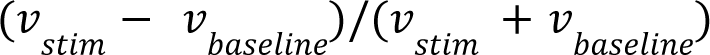

Where 𝑣_𝑠𝑡𝑖𝑚_ is the average jaw velocity during the first 5 seconds of stimulation, and 𝑣_𝑏𝑎𝑠𝑒𝑙𝑖𝑛𝑒_ is the average jaw velocity during the 5 seconds before stimulation. The change in jaw velocity was averaged across trials for a given stimulation pattern. Significance was determined by whether this average exceeded the 99% confidence interval of a bootstrap distribution. The bootstrap distribution was estimated by randomly selecting new timepoints to calculate the above average using 10,000 times. Error bars show the standard error of the mean. To determine the effect of mouse group (opto vs control, Figure 6D), we performed a two sample t-test on the change in jaw velocity. To determine the effect of mouse group and stimulation location, we performed a two-way interaction ANOVA on the change in jaw velocity.

## RESOURCE AVAILABILITY

Our pipeline and MATLAB code for fiber localization from CT images can be found here: (https://github.com/HoweLab/MultifiberLocalization). The smaller diameter of the fibers makes them more flexible than larger fibers, and as a result, implantation may change the spatial relationship between the tips and some fibers may even end up crossing (Figure 1E). It is for this reason that we rely on the 3D CT scan of the implanted brain for unambiguous localization of these densely arranged fibers. Nonetheless, it may be possible for those without access to a micro-CT scanner to localize fibers using more conventional histological methods. In order to accurately localize the tip, users might need to follow the fiber track across multiple slices. A1potential avenue might be to reconstruct the implanted brain volume from sequential slices into a 3D image, and then adapt our code to localize fibers in the reconstructed 3D image.

## ACKNOWLEDGMENTS

This work was supported by the following funding sources:

MWH - Aligning Science Across Parkinson’s (PF-SF-JFA-836662), National Institute of Mental Health (R01 MH125835), Whitehall Foundation Fellowship, Klingenstein-Simons Foundation Fellowship, Parkinson’s Foundation (Stanley Fahn Junior Faculty Award); MTV - NIMH F32MH120894; EHB - NIMH 1F31NS127536; KJM - NIDCD F32DC020665; IGD – NINDS RF1NS128975. We thank the Boston University Centers for Neurophotonics and Systems Neuroscience for financial and technical support, Micro CT Core for providing equipment and technical expertise for micro-CT scanning, and Boston University Animal Science Center for providing central laboratory animal care and support resources.

## AUTHOR CONTRIBUTIONS

Conceptualization - MWH, DAB, MTV, EHB; Methodology - MTV, EHB, MJW, CAN, ZZ, KJM, SB, LM, TMO, IGD, DAB, MWH; Software - MTV, BMG; Formal Analysis - MTV, EHB, CAN, ZZ, KJM, SB, BMG, MWH; Investigation - MTV, EHB, MJW, CAN, ZZ, KJM, SB, LM; Resources - BMG, YZ, YL, TMO; Data Curation - MTV, EHB; Writing Original Draft - MTV, EHB, ZZ, KJM, SB, IGD, DAB, MWH; Visualization - MTV, EHB, CAN, ZZ, KJM, SB, IGD, MWH; Supervision - IGD, DAB, MWH; Funding Acquisition - MTV, EHB, KHM, IGD, DAB, MWH.

## INCLUSION AND DIVERSITY STATEMENT

We work to ensure sex balance in the selection of non-human subjects. One or more of the authors of this paper self-identifies as an underrepresented ethnic minority in their field of research or within their geographical location. One or more of the authors of this paper self-identifies as a gender minority in their field of research. One or more of the authors of this paper self-identifies as a member of the LGBTQIA+ community. We support inclusive, diverse, and equitable conduct of research.

## SUPPLEMENTAL INFORMATION

**Supplemental Figure 1:**
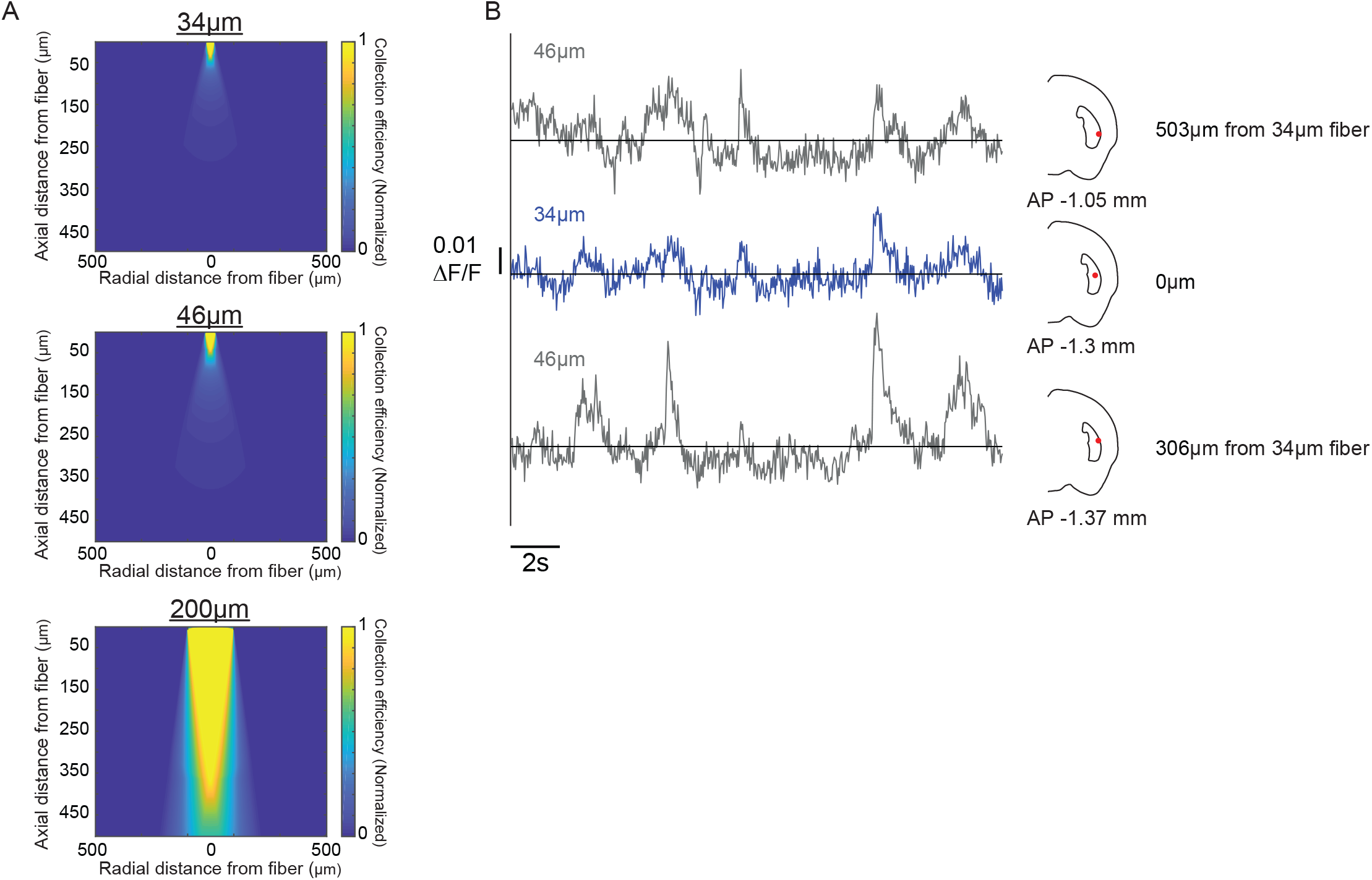
Collection efficiency and signal through small diameter optical fibers. **(A)** Collection efficiency estimated at radial and axial distances from the tips of two types of optical fibers (46µm core and 34µm core, 0.66NA) tested with the multi-fiber arrays and a 200µm (0.37NA) core fiber used commonly in conventional photometry experiments^15,66^. Smaller diameter fibers enabled more restricted sampling and our spacing ensured very minimal overlap between collection fields of neighboring fibers. **(B)** dLight1.3b ΔF/F traces from two representative 50µm diameter fibers (gray) and one 37µm diameter fiber (blue) located in close proximity in the brain (locations indicated on right).

**Supplemental Figure 2:**
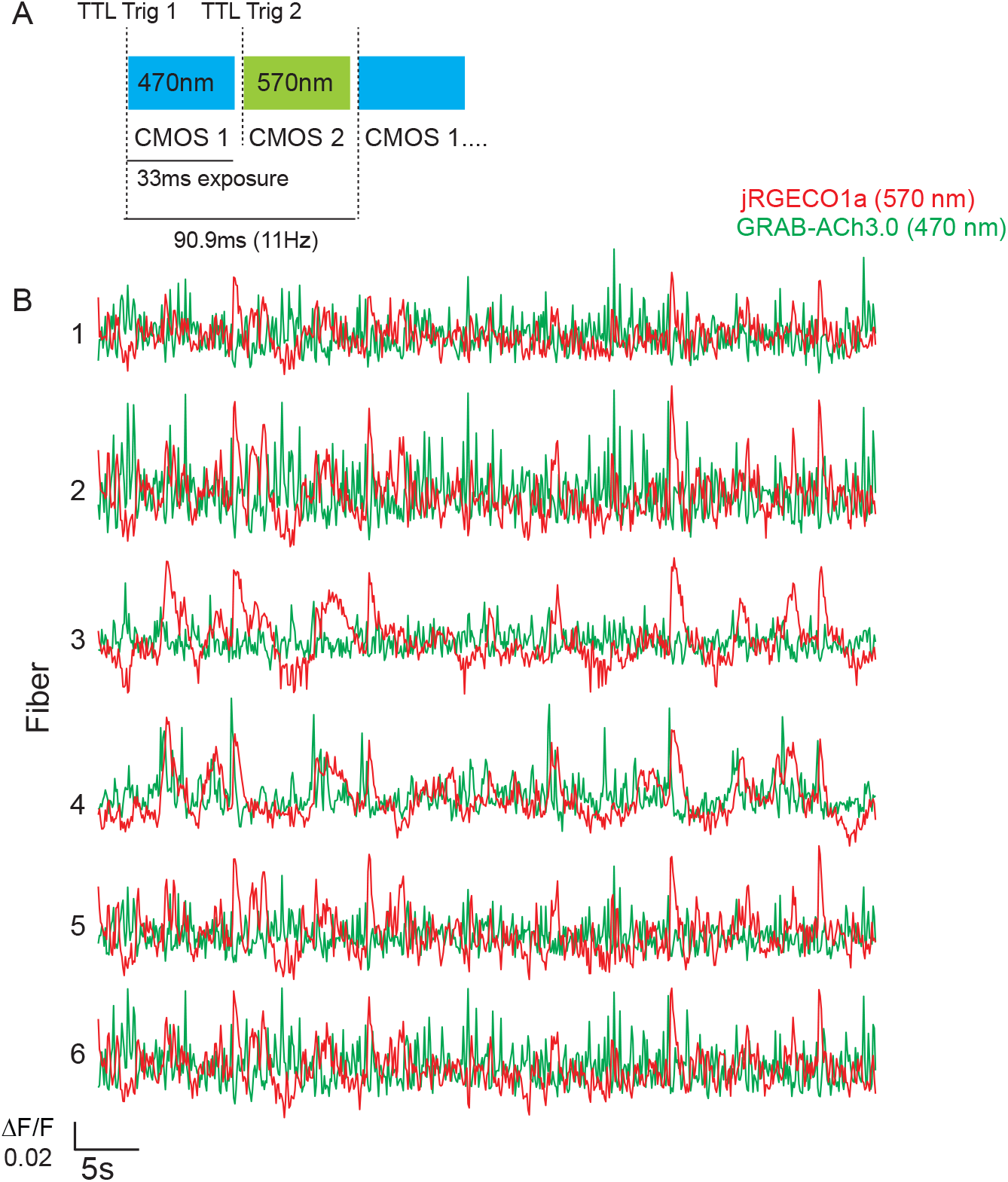
Quasi-simultaneous imaging of red and green fluorescent indicators through multi-fiber arrays. **(A)** Setup for dual-wavelength imaging. TTL pulses (5V) from a NIDAQ simultaneously triggered LEDs to excite green and red fluorophores (470nm and 570nm respectively) and the corresponding CMOS camera in the green and red collection path (see Figure 1). Pulses to each LED/camera were alternated at 22Hz to achieve a frame rate of 11Hz and a 33ms exposure time for each camera. **(B)** Quasi-simultaneous measurements of red and green fluorescent indicators for calcium and acetylcholine. Representative ΔF/F traces from six 50µm fibers in a multi-fiber array recorded from the striatum of a head-fixed, behaving mouse expressing jRGECO1a (red) in axons of midbrain DA neurons and GRAB-Ach3.0 (green) in striatal neurons.

**Supplemental Figure 3:**
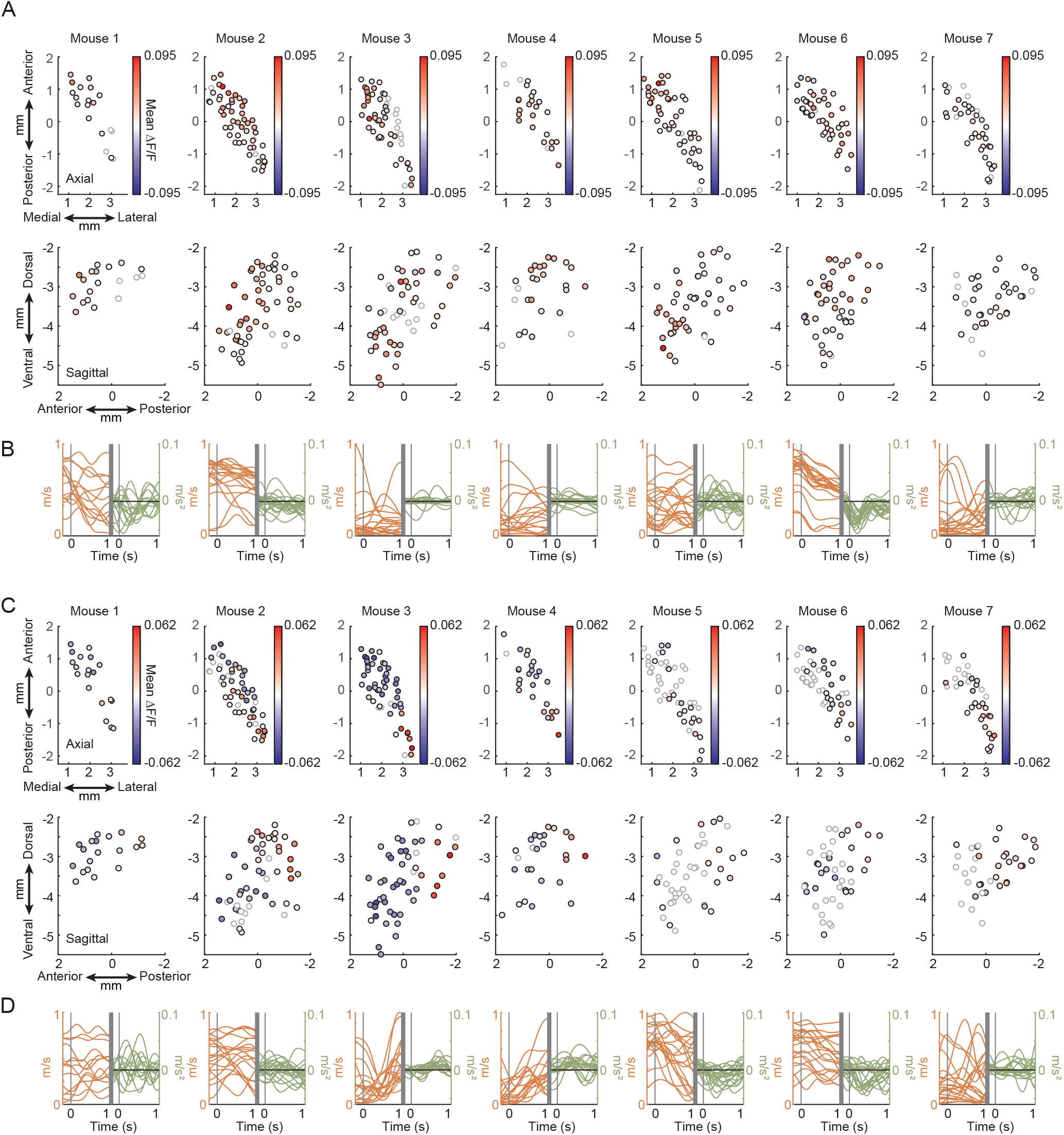
DA release patterns to salient stimuli in individual mice. **(A)** Largest amplitude mean ΔF/F change across all light presentations for individual mice (n = 7, same mice as Figures 4, 5) in a 1-second window after stimulus onset. The color of each circle represents the largest mean ΔF/F change for a single fiber. The location of each circle indicates its localized fiber position in the axial (top) and sagittal (bottom) planes. Open circles, non-significant increases or decreases relative to baseline (mean exceeding 99% confidence interval of the bootstrapped null distribution for 3 consecutive timepoints, p<1.0×10^-6^). **(B)** Total velocity (orange, left) and acceleration (green, right) aligned on light onset for all light presentations for each mouse indicated in **A**. **(C)** Same as **A** but for tone presentations. **(D)** Same as **B** but for tone presentations.

**Supplemental Figure 4:**
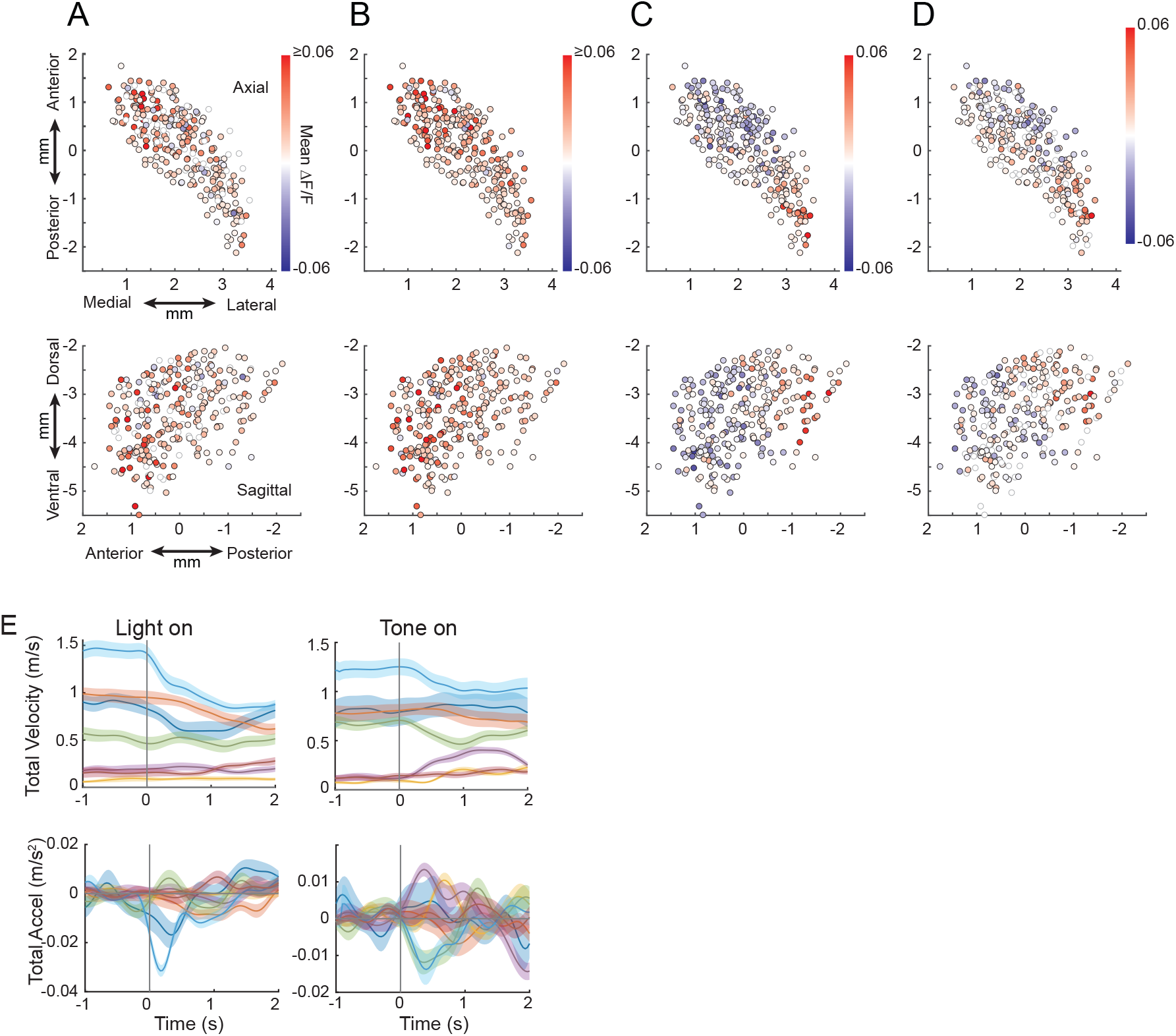
Motor reactions to salient stimuli cannot account for modality specific patterns of DA release. **(A)** Largest amplitude mean ΔF/F change across all light presentations that were followed by a locomotor acceleration for all fibers (n = 280) and mice (n = 7, same mice as Figures 4, 5) in a 1-second window after stimulus onset. The color of each circle represents the largest mean ΔF/F change for a single fiber. The location of each circle indicates its localized fiber position in the axial (top) and sagittal (bottom) planes. Open circles, non-significant increases or decreases relative to baseline (bootstrap test vs null distribution for 3 consecutive timepoints, p<1.0×10^-6^). **(B)** Same as **A** but for presentations followed by a deceleration. **(C)** Same as **A** but for tone presentations followed by locomotor accelerations. **(D)** Same as **C** but for tone presentations followed by a deceleration. **(E)** Mean velocity (top) and acceleration (bottom) averaged across all light (left) and tone (right) presentations for each mouse. Each color is the mean trace from a single mouse. Shaded regions, SEM.

**Supplemental Figure 5:**
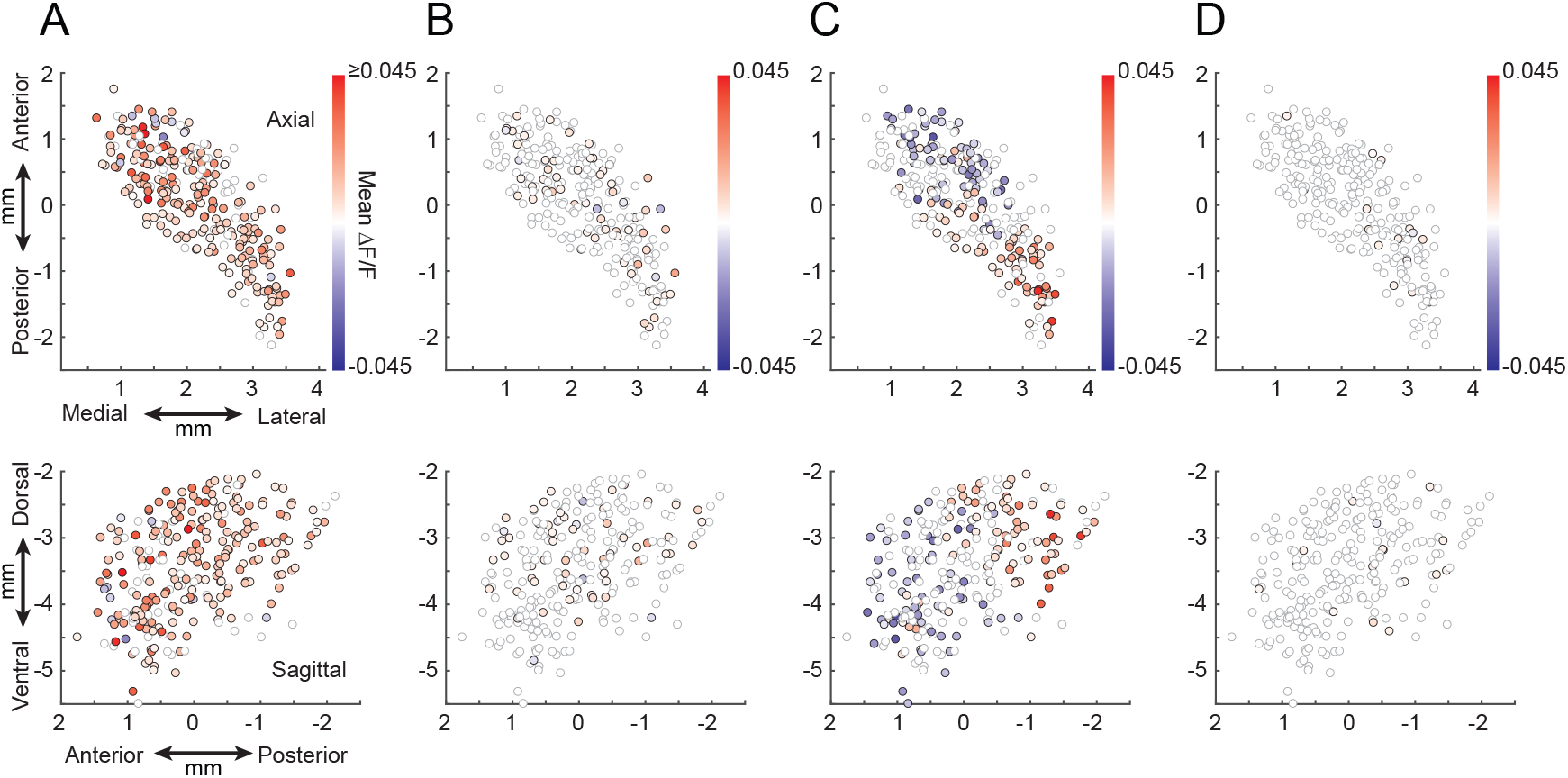
Hemodynamic and motion artifacts do not contribute significantly to salient stimulus evoked DA release patterns. **(A)** Largest amplitude mean ΔF/F change for 470nm excitation in a 1-second window after all light presentations (n = 10 trials) during a quasi-simultaneous isosbestic control session where 470nm excitation light was alternated with 405nm light at 36Hz, as in Figure 1C. Data is shown for 261 fibers from 6/7 of the mice from Figures 4, 5 (1 mouse did not have data form an isosbestic control session). The color of each circle represents the largest mean ΔF/F change for a single fiber. The location of each circle indicates its localized fiber position in the axial (top) and sagittal (bottom) planes. Open circles, non-significant increases or decreases relative to baseline (bootstrap test vs null distribution for 3 consecutive timepoints, p<1.0×10^-6^). **(B)** Same as **A** but for quasi-simultaneous 405nm excitation. **(C)** Same as **A** (470nm excitation) but for tone presentations (n = 10 trials) during the quasi-simultaneous isosbestic control session. **(D)** Same as **C** but for 405nm excitation.

**Supplemental Figure 6:**
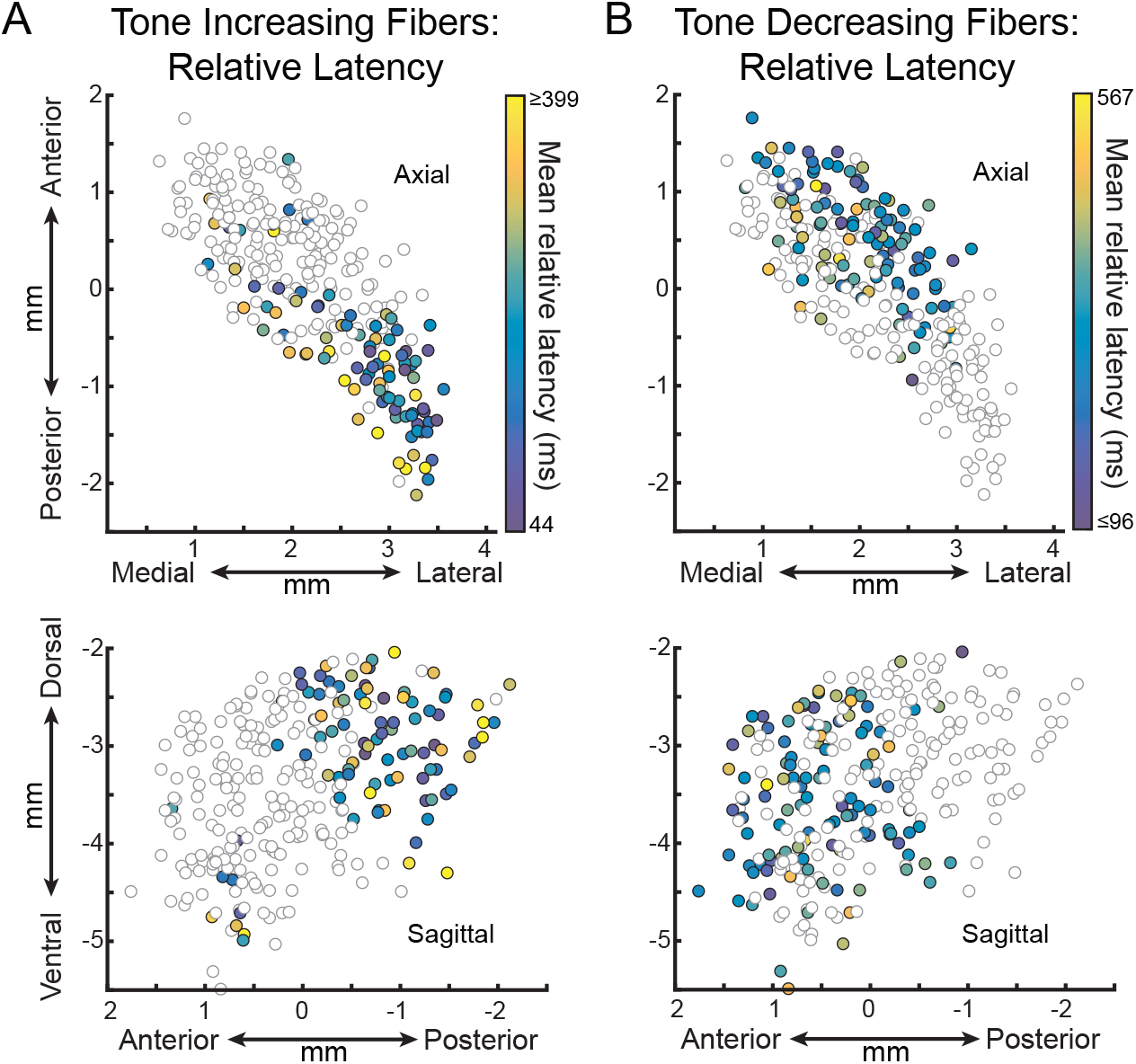
Spatial distribution of latencies for increases and decreases in DA release to tone presentations. **(A and B)** Axial (top) and sagittal (bottom) spatial maps of the mean relative half-max (A) or half-min (B) latencies of DA release for each fiber across all tone presentations for all fibers (n = 280) and mice (n = 7, 4F + 3M). The color of each circle represents the mean relative latency for a single fiber. Open circles in spatial maps are non-significant increases or decreases relative to baseline (mean triggered average exceeding 99% confidence interval of the bootstrapped null distribution for 3 consecutive timepoints, p<1.0×10^-6^). (A) displays half-max latencies for fibers with significant increases to the tone and (B) displays half-min latencies for fibers with significant decreases. Normalization is relative to trial-by-trial latencies for fibers with significant increases or decreases for (A) and (B) respectively.

**Supplemental Figure 7:**
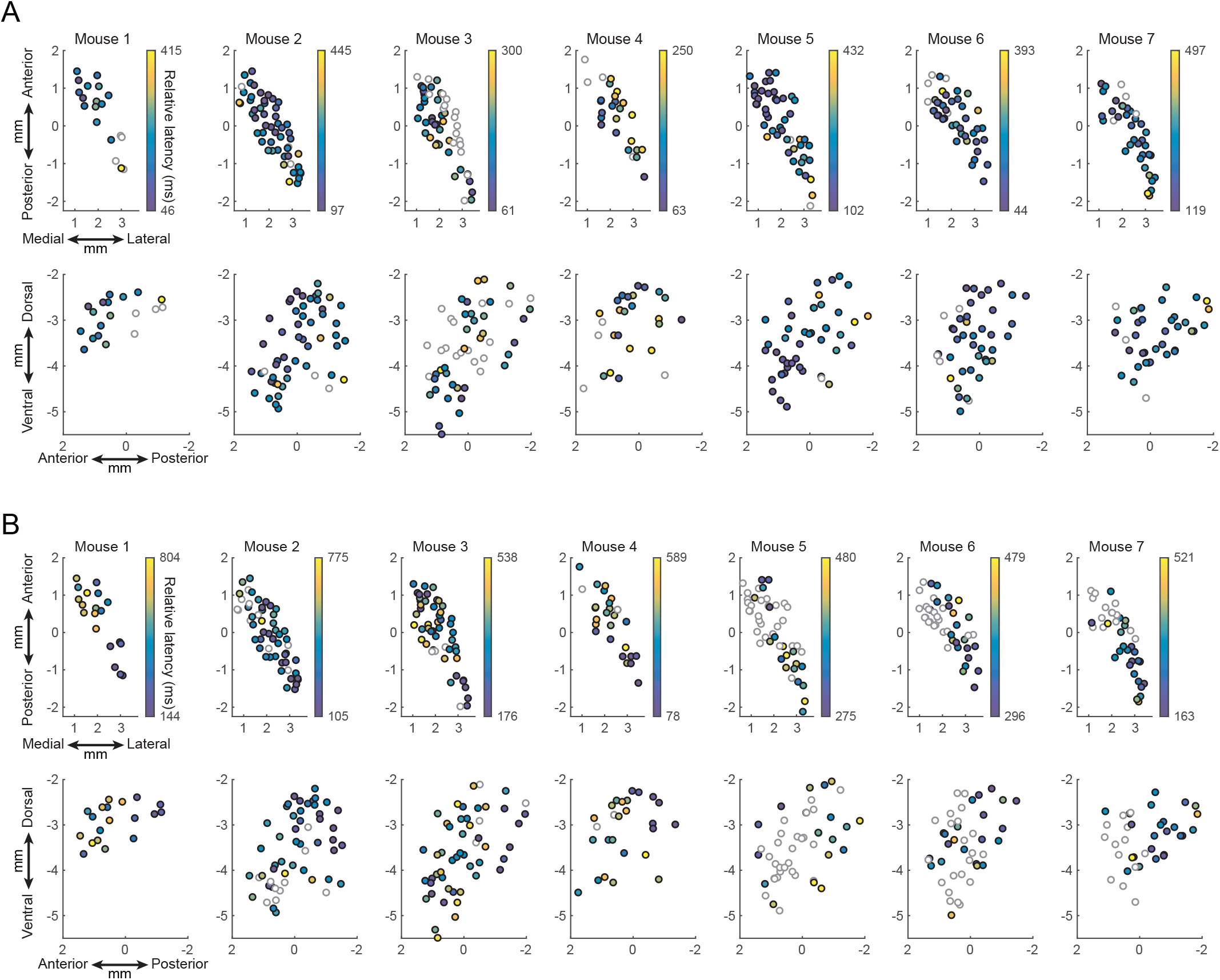
Spatial distribution of DA release latencies to salient stimuli in individual mice. **(A)** Mean relative latencies of DA responses as in Figure 5 for all fibers in individual mice (n = 7, same mice as Figures 4, 5). **(B)** Same as A, for tone presentations.

Supplemental Table 1.

**Supplemental Table 1**
| Construct | Concentration (GC/mL) | Source | Purpose | Experiment(s) |
| --- | --- | --- | --- | --- |
| pAAV-CAG-dLight1.3b (AAV5) | $3.75 \times 10^{12}$ - $6.00 \times 10^{12}$ | Addgene | green dopamine sensor | Fig. 1d<br>Fig. 2c<br>Fig. 4<br>Fig. 5<br>Supp 1<br>Supp 3<br>Supp 4<br>Supp 5<br>Supp 6<br>Supp 7 |
| pGP-AAV-syn-FLEX-jGCaMP7f-WPRE | $3.60 \times 10^{12}$ - $7.00 \times 10^{12}$ | Addgene | green calcium sensor | Fig. 1c<br>2a-b |
| AAV9-hSyn-Ach3.0 | $7.65 \times 10^{12}$ | WZ Biosciences | green acetylcholine sensor | Supp 2 |
| pAAV-syn-FLEX-NES-jRGECO1a-WPRE-SV40 (AAV1) | $1.47 \times 10^{13}$ | Addgene | red calcium sensor | Supp 2 |
| pAAV-EF1a-double floxed hChR2(H134R)EYFP-WPRE-HGHpA | $1.05 \times 10^{13}$ | Addgene | Blue light opsin | Fig. 6<br>Fig. 7 |
| pAAV-hsyn-rDA3m (AAV 2/9) | $8.79 \times 10^{10}$ | biohippo | Red dopamine sensor | Fig. 6 |

